# Coordinated tri-inhibition generates human CA1, CA3 and DG neurons and hippocampal astrocytes forming long-lived and electrophysiologically complex cultures

**DOI:** 10.64898/2026.01.21.700748

**Authors:** Kwaku Dad Abu-Bonsrah, Candice Desouza, Hui Wen Chan, Forough Habibollahi, Brad Watmuff, Mirella Dottori, Brett J. Kagan

## Abstract

The diverse computational functions of the human hippocampus rely on coordinated interactions among dentate gyrus (DG), CA3, and CA1 subfields, yet generating all three neuronal identities in vitro - particularly CA1 - has remained challenging. Here we establish a reproducible and modular differentiation protocol that directs human pluripotent stem cells through dorsomedial telencephalic progenitors to DG, CA3 and CA1 neurons and regionally specified hippocampal astrocytes. Early tri-inhibition with sonic hedgehog inhibition generated FOXG1+PAX6+ forebrain progenitors, while FGF2 treatment maintained the progenitors and induced TBR2+ intermediate progenitors, and controlled WNT activation with CHIR99021 enriched PROX1+ hippocampal progenitors in two donor lines. Bulk RNA-seq confirmed hippocampal identity, and a promoter-driven reporter for a CA1-specific gene [*MPPED1*] validated CA1 neurons, alongside DG (PROX1), CA3 (ELAVL2, GRIK4, SCGN) and CA1 (WFS1, OCT6) markers and maturing synaptic puncta. Hippocampal organoids remained active for more than 400 days, with higher firing rates, information entropy and hub-like causal connectivity than monolayers; astrocyte-supplemented monolayers showed intermediate maturation. This platform enables systematic experimental control of neuron-astrocyte ratio, culture geometry and developmental timing for studying human hippocampal development, circuit function and disease.

## Introduction

The hippocampus functions as a dynamic computational hub for cognition, memory, and adaptive behaviour by integrating multimodal information and updating representations based on context and experience^1^. Through interactions with cortical and subcortical regions, it supports relational and sequential memory, spatial mapping, and flexible inference, mediated by specialised circuits for pattern completion and unique networks of place and grid cells for spatio-temporal coding^1–3^. These processes rely on balanced excitatory–inhibitory activity and plasticity mechanisms, including long-term potentiation (LTP) and long-term depression (LTD), which underpin learning and memory consolidation^4–6^. Theoretical frameworks – from declarative and relational models to spatial navigation theories – highlight its central role in episodic and semantic memory, as well as predictive modelling^1^. Moreover, adult hippocampal neurogenesis is one of the few sources of new neurons in adult humans and has been recognised as a key contributor to brain plasticity and cognitive resilience^7,8^. To understand these numerous essential functions, elucidating hippocampal cellular independent function, along with network interactions of the hippocampus within the wider brain, is essential.

In humans, the hippocampus sits as a C-shaped structure in the medial temporal lobe which originates from the dorsomedial telencephalon^9,10^. It comprises the dentate gyrus (DG), cornu ammonis (CA1–CA4), and subiculum, which together form the trisynaptic circuit for memory processing. From the cornu ammonis, CA1 and CA3 have been identified as particularly critical for cognitive tasks^4,6,11–13^. CA3 drives pattern completion and associative memory, CA1 integrates and outputs processed information, while the dentate gyrus and subiculum mediate cortical input and memory consolidation^14–17^. The complexity of hippocampal function necessitates in vitro models to study its computational processes under controlled conditions. These models allow detailed investigation of spatial and temporal task performance, synaptic and network dynamics, and mechanisms underlying plasticity, memory formation, and disease. With the discovery of human induced pluripotent stem cells (hPSCs), scalable and ethical generation of living region-specific neurons has become possible^18^. Several protocols have been previously reported for generating hPSC-derived neurons based on a variety of different methods^19–23^. Neural induction through the embryoid body (EB) formation is one of the most common approaches^19^. Another widely implemented method for neural induction is inhibition of the transforming growth factor-β/SMAD signaling pathway by Noggin/LDN193189 and SB431542^24^. However, while these protocols often generate CA3 and DG neurons, CA1 neurons have been identified as a critical missing cell type from these methods^25,26^.

To develop a differentiation protocol that was able to produce the full range of cell types of interest, we considered the ontogenetic development of the hippocampus. The hippocampus originates from the medial pallium structures of the dorsomedial telencephalon. The dorsomedial telencephalon consists of the choroid plexus, the cortical hem, and the medial pallium. The cortical hem lies between the choroid plexus and the medial pallium structures of the dorsomedial telencephalon. The cortical hem serves as a source of signaling cues secreting molecules, Wingless/Integrated (Wnt) and Bone Morphogenetic Protein (BMP). Wnt signaling, particularly via Wnt3a, plays a crucial role in initiating and organising early hippocampal formation and neurogenesis through downstream transcription factors at the caudomedial margin of the continuous cerebral cortical neuroepithelium^27–30^. In mice and humans, the role of Wnt3a in hippocampal specification has been highlighted^9,22^. There are specific molecular markers that define hippocampal subfields. The gene *GRIK4* (glutamate receptor, ionotropic, kainate 4), which encodes the *KA1* kainate receptor subunit, is enriched in the CA3 region^31^, while Wolfram syndrome 1 (*WFS1*) and POU domain, class 3, transcription factor 1 (*POU3F1*, also known as *OCT6*) expression is enriched in CA1^10,32^. The gene, Prospero Homeobox 1 (*PROX1*) is also enriched in the DG. Finally, in hippocampal development and growth, Fibroblast growth factor (FGF) signaling has a multifactorial role-proliferation and survival^33,34^.

Beyond only focusing on neurons however, hippocampal region-specific astrocytes were also considered a key target from the differentiation effort. Critical region-specific functional differences between astrocytes have also been observed, highlighting the need to be able to generate not just region-specific neurons, but also astrocytes^35–38^. Astrocytes are the most abundant cell type in the brain, and their function was initially thought to support the neurons passively^39^. However, current findings have found the role of astrocytes in the brain and their function in both neurodevelopmental and neurodegenerative diseases to be significantly more complex^40^. As an example, astrocytes also play a key role in neuronal precursor guidance and maturation of neurons^41^. Supporting this, coculturing of neurons and astrocytes substantially improves the functional maturation of human pluripotent stem cell-derived neurons^42^. Alternatively, astrocytes and neurons can be differentiated from a common neural progenitor, giving rise to the two different cell types. However, the use of this method makes it difficult to monitor and control the exact percentage of astrocytes, and limits establishing different ratios of neurons to astrocytes, or even culturing only neurons as an experimental control^43,44^. For this reason, we also explored adapting our protocol to be able to develop region-specific astrocytes from the hippocampal lineage.

Ultimately, this work aimed to develop and validate a new method for generating fully specified hippocampal differentiation, with a particular focus on CA1, CA3, and DG neurons, along with hippocampal regionally specified astrocytes. This work was validated using immunocytochemistry and qPCR on hPSCs from two donors to capture both XX and XY chromosomal karyotypes. By using hPSCs from two donors and testing across both monolayers and organoids we were able to show the stability of this approach. Consequently, we also performed long-term electrophysiological recordings of each culture structure and tracked the differing maturation and properties of these cultures over time, including periods over a year for some. Importantly, we validated the generation of CA1 neurons using an *MPPED1* promoter-driven reporter strategy, providing independent confirmation of CA1 identity and demonstrating the capacity of this differentiation protocol to robustly generate all three major hippocampal subfield populations.

## Results

### Efficient generation of hippocampal progenitors from hPSCs via WNT activation to induce PROX1+ hippocampal progenitors

Hippocampal progenitors are known to originate from the medial pallium as directed by the signal molecules from the cortical hem^10^. There are established differentiation protocols to induce neocortex, also known as cortical neuroepithelium (NE). Following a previously described neocortex induction method^45,46^ (**Figure 1A**), we induced hPSCs into dorsomedial telencephalic tissues using the tri-inhibition (dual SMAD and Wnt inhibition) protocol. This has been shown to induce PAX6+ cortical NE in culture^46,47^. After 2 days of induction, we further blocked ventralisation by using cyclopamine, which inhibits the sonic hedgehog (SHH) signalling pathway^48,49^. The cultures at day 11 showed readily observable rosette structures validating the right specification of the differentiation protocol (**Figure 1B**). To investigate the expression of *EMX2*^50^ and *GLI3*^51^, which are expressed in the dorsomedial telencephalon, as well as the proneural genes *PAX6* and *DCX*, expressed in embryonic hippocampal progenitors^28,52^, we analysed the RNA expression at day 11 in vitro (DIV 11). Transcriptional analysis showed the increased levels of *EMX2, PAX6, DCX* and *GLI3* (**Figure 1C**). The differentiation robustly induced PAX6+ dorsal progenitors and OTX2+ forebrain progenitor (95 ± 2%) with 98 ± 1% of these cells expressing the gene *NESTIN*, a marker for neural stem cells/progenitor cells^53^ (**Figure 1D & E**). The gene *FOXG1* is a telencephalic marker, expressed from medial pallium to neocortex, and we used *FOXG1* to gauge the development of neuroprogenitors. We noted 96 ± 0.5% induction of FOXG1+ progenitors (**Figure 1D & E**). These observations, together with the immunocytochemistry data, demonstrate the induction of neural progenitors in the dorsomedial telencephalon region.

**Figure 1:**
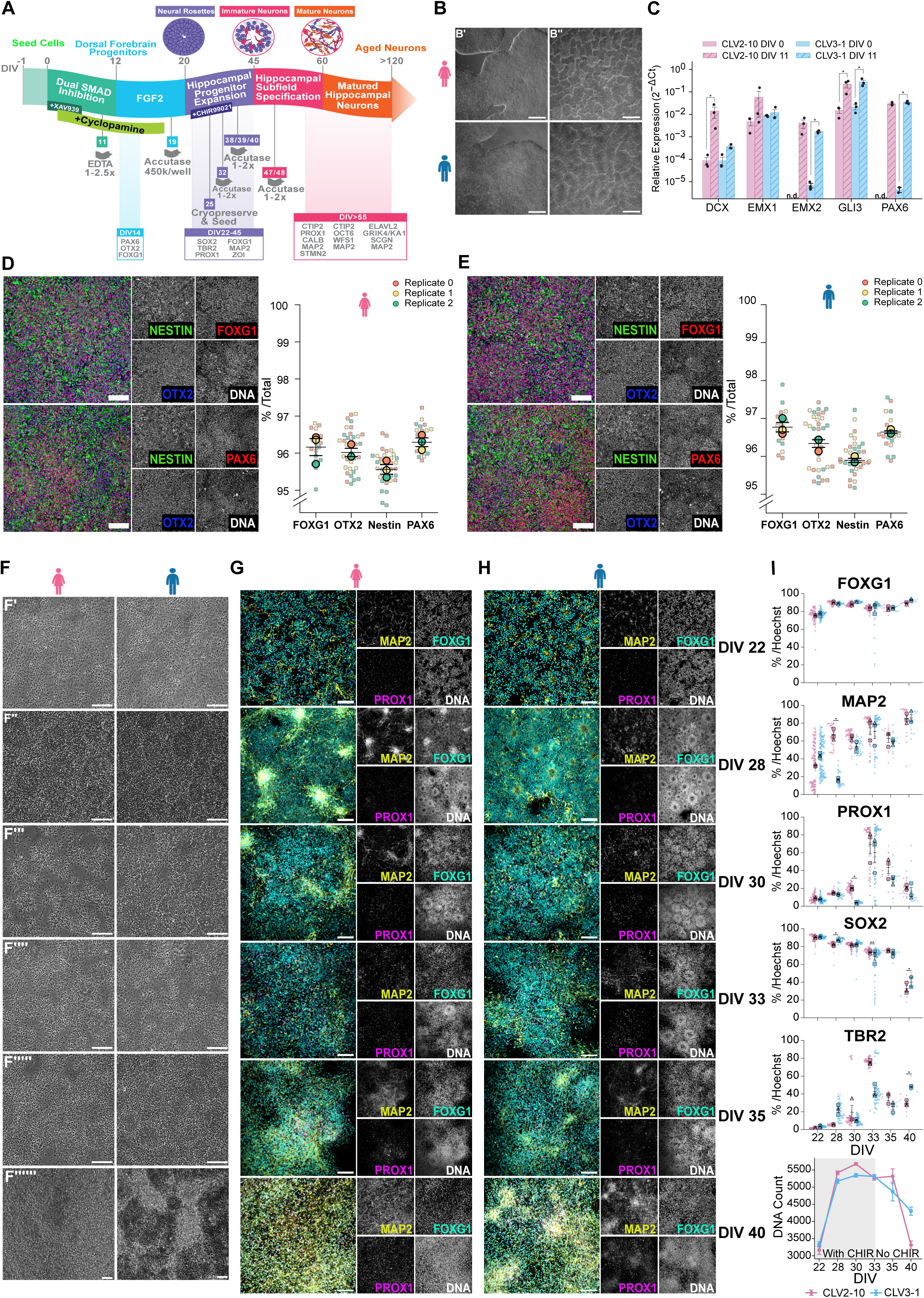
Efficient generation of TBR2+ IPCs and PROX1+ neurons from dorsomedial telencephalic-like tissues. A- Schematic representation of the differentiation protocol showing tri-inhibitory inhibition and SHH inhibition to generate dorsomedial telencephalic progenitors from hPSCs. Progenitors are further differentiated and matured till further analysis. B- Representative brightfield images of stem cells (B’) and differentiated cells at DIV 11 (B’’). C- qRT-PCR confirms the expression of dorsomedial telencephalic markers. Barplots with no data points indicate that no measurements were obtained for that marker on the given day in vitro (n = 3 independent experiments each, data are mean ± SEM). The Y axis refers to the % of cells expressing the marker over the total of cells counted as per Hoechst staining. D- CLV2-10 (female) hPSCs-derived hippocampal progenitors at DIV 14 express dorsomedial telencephalic markers OTX2, PAX6 and FOXG1 and the neural stem cell marker, NESTIN. Quantification of FOXG1, OTX2, PAX6 and NESTIN at DIV 14 (n = 3 independent experiments each, data are mean ± SEM). The Y axis refers to the % of cells expressing the marker over the total of cells counted as per Hoechst staining. Scale bar is 100 µm. E- CLV3-1 (male) hPSCs-derived hippocampal progenitors at DIV 14 express dorsomedial telencephalic markers OTX2, PAX6 and FOXG1 and the neural stem cell marker, NESTIN. Quantification of FOXG1, OTX2, PAX6 and NESTIN at DIV 14 (n = 3 independent experiments each, data are mean ± SEM). The Y axis refers to the % of cells expressing the marker over the total of cells counted as per Hoechst staining. Scale bar is 100 µm. F- Representative brightfield images of differentiated hPSCs-derived hippocampal progenitors from DIV 22-40 (F’-F’’’’’’). Scale bar is 500 µm G- Representative immunostain of the female hPSCs-derived hippocampal progenitors showing the appearance of PROX1+ cells from DIV 22-40. Scale bar is 100 µm. H- Representative immunostain of the male hPSCs-derived hippocampal progenitors showing the appearance of PROX1+ cells from DIV 22-40. Scale bar is 100 µm. I- Time-course quantification of FOXG1⁺ and SOX2⁺ progenitor populations, TBR2+ IPCs and PROX1+ progenitor populations and MAP2⁺ neuronal cells for two hPSCs. CHIR99021 treatment increased total number of cells in culture for the two hPSC lines (male and female n = 3 independent experiments each, data are mean ± SEM). Data are mean ± SEM, with confidence intervals defined by the minimum and maximum observed values. The Y axis refers to the % of cells expressing the marker over the total of cells counted as per Hoechst staining. * *p* < 0.05, ** *p* < 0.01, \*\*\**p* < 0.001

We expanded the neural progenitors by treating the cultures with FGF2 for 8 days based on previously published findings^45^ (**Figure 1A**). This promoted the generation of SOX2+ and FOXG1+ neural progenitor cells (NPCs) and TBR2 (also known as EOMES)-positive intermediate progenitor cells (IPCs). The gene TBR2 is a transcription factor regulating both postnatal and adult hippocampal neurogenesis. In rodent models, TBR2 expression has been shown to modulate IPC proliferation and drive PROX1 expression^54^. At DIV 22, immunostaining confirmed the presence of TBR2+ IPCs (5 ± 1%) with the SOX2+ cells self-organising into rosettes with ZO-I+ lumens (**Supplementary Figure 1A** & **B**). Transcriptional analysis of DIV 19-30 confirmed the increased expression of pro-neural genes (notably *PAX6, ASCL1*), embryonic hippocampal progenitor genes (notably *PROX1* and *DCX*), and downregulation of the pluripotency gene, *OCT4* (**Supplementary Figure 2A**).

Wnt signaling regulates proliferative or neurogenic fate decisions in hippocampal progenitors and has been shown to play a key role in hippocampal neurogenesis. Wnt signaling regulates *PROX1* expression, a marker essential for DG cell fate and adult hippocampal neurogenesis^23,55^. The classic GSK-3β inhibitor (Wnt agonist), CHIR99021, activates the canonical Wnt signaling pathway and upregulates the Wnt genes expressed in the cortical hem. Recent findings have shown that treating NPCs with CHIR99021 will induce neural progenitor cell proliferation and further differentiation of IPCs into PROX1+ cells^28,30^. We next sought to investigate the impact of CHIR99021 treatment on number of cells as WNT signalling regulates cell proliferation and is essential in NPC proliferation in the medial pallium^30^. We first tested two different concentrations of CHIR99021 (1.5 µM and 3 µM), and we noted the slight increase in number of nuclear positive (Hoechst+) cells in the 3 µM-treated cells as compared to 1.5 µM-treated groups. Immunostaining for PROX1 showed both conditions induced the differentiation of PROX1+ cells, with treatment of 3 µM CHIR99021 leading to increased expression of PROX1+ and TBR2+ cells as compared to 1.5 µM at DIV 40. We then decided to go ahead with 3 µM as previously published^9,56^, and monitor the sturdy increase in PROX1 expression (**Supplementary Figure 2B**).

We found that PROX1 expression increased steadily from DIV 28-40. PROX1 expression was detected at DIV 22, and its expression increased steadily after CHIR99021 treatment. As expected, the TBR2 and PROX1 expression increased steadily, which confirms previous findings in rodents^54^. The expressions of FOXG1 and SOX2 remained unchanged over the days till DIV 35, suggesting the constant differentiation ability of the cells. The decline in SOX2+ progenitors, together with the corresponding increase in MAP2+ cells, indicates that the neurons have begun to undergo terminal differentiation. The steady increase of both TBR2+ IPCs and PROX1+ cells indicates the proliferation and differentiation ability of the TBR2+ IPCs, giving rise to new PROX1+ and other neural progenitors (**Figure 1F, G, H & I**). Transcriptional analysis of cells at DIV 19-30 confirmed the induction of genes expressed in both hippocampal progenitors and differentiating cells (**Supplementary Figure 2A**). We confirmed the proliferative response to CHIR99021 by quantifying total cell numbers from DIV 22-40, during which cultures showed a ∼1.5-fold increase before plateauing. These findings indicate that CHIR99021 enhances cell proliferation and supports the generation of hippocampal progenitors by increasing the production of PROX1+ neurons, with no detectable differences across the hPSC lines examined.

### Generation of CA1 and CA3 hippocampal pyramidal- and granule-like neurons

To evaluate whether terminally differentiated neural progenitor cells give rise to hippocampal-specific neuronal subtypes, we analysed the expression of genes associated with the dentate gyrus (DG), CA1, and CA3 subfields^57^ (**Figure 2A**). We first confirmed the increased expression of pro-neural genes (notably *PAX6, ASCL1*), embryonic hippocampal progenitor genes (notably *PROX1* and *DCX*) and neuronal genes (notably *MAP2*, *NEUN*, *SYN1*) by performing transcriptional analysis of DIV 37-54 (**Supplementary Figure 2A**). Mature neurons are characterised by expression of microtubule-associated proteins (MAPs), which regulate cytoskeletal stability; TAU is the predominant MAP in mature neurons^58^. In our cultures, MAP2 was used to label dendrites and TAU to identify axons and assess neuronal maturation. Immunostaining after DIV 60 confirmed the presence of 60 ± 3% MAP2+ cells (**Supplementary Figure 3A**).

**Figure 2:**
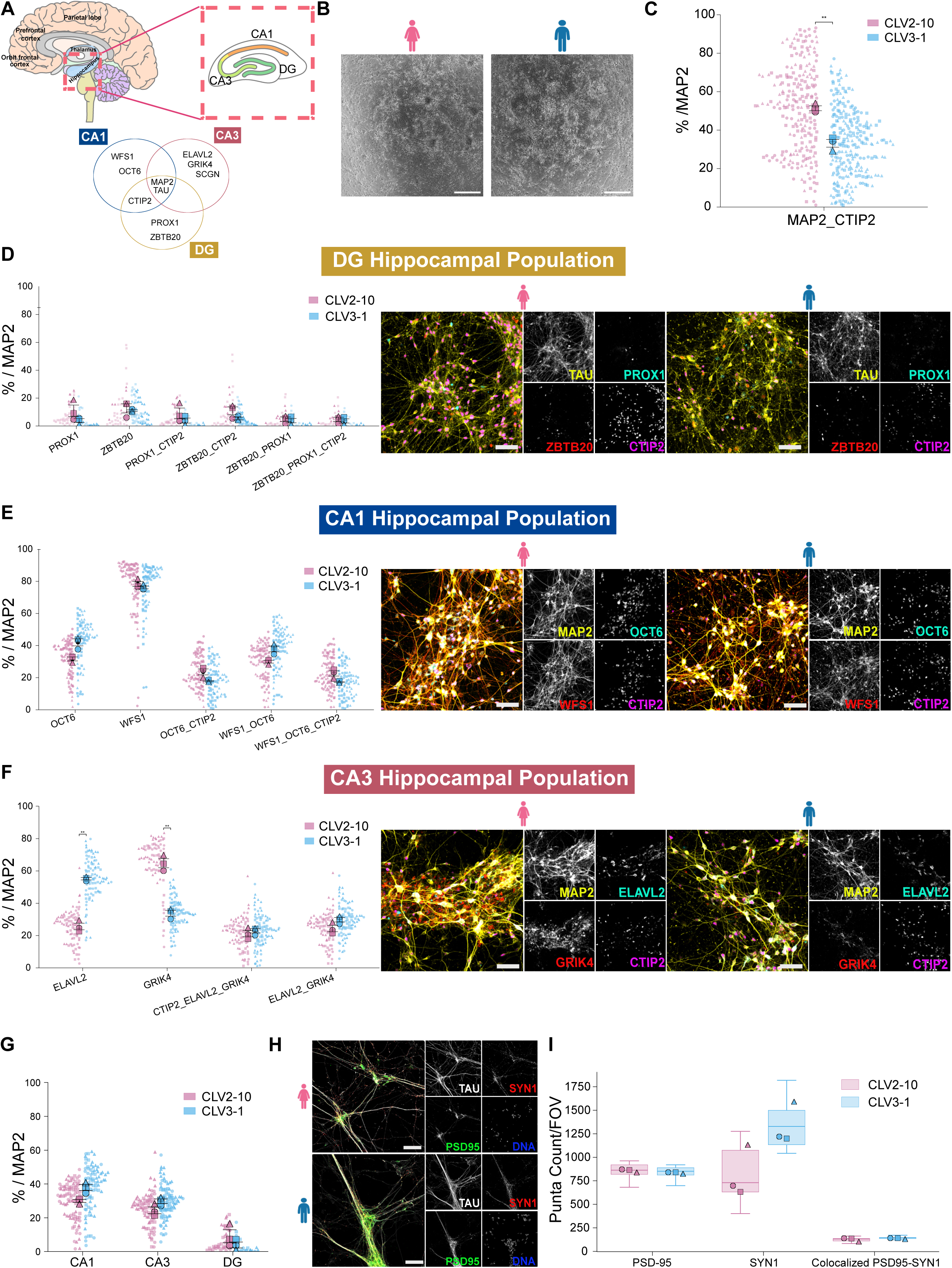
Generation of neurons constituting the different hippocampal subfields (DG, CA1 and CA3) upon long term culture (DIV 90). A- Schematic of the human hippocampus and its major subfields within a sagittal brain section. The Venn diagram illustrates subfield-specific and overlapping gene expression signatures. B- Representative brightfield images of differentiated hPSCs-derived hippocampal progenitors after long-term culture. Scale bar is 500 µm C- Quantification of CTIP2+ cells in total MAP2+ cell population of the two hPSC lines (male and female n = 3 independent experiments each, data are mean ± SEM). D- Immunostaining and quantification analysis of ZBTB20+, and PROX1+ cells in total TAU+ cell population constituting distinct DG hippocampal neurons for two hPSC lines (male and female n = 3 independent experiments each, data are mean ± SEM). Scale bar is 100 µm. E- Immunostaining and quantification analysis of OCT6+ and WFS1+cells in total MAP2+ cell population constituting distinct CA1 hippocampal neurons for two hPSC lines (male and female n = 3 independent experiments each, data are mean ± SEM). Scale bar is 100 µm. F- Immunostaining and quantification analysis of ELAVL2+ and GRIK4+cells in total MAP2+ cell population constituting distinct CA3 hippocampal neurons for two hPSC lines (male and female n = 3 independent experiments each, data are mean ± SEM). Scale bar is 100 µm. G- Quantification of the proportion of the different hippocampal subfield population in the differentiation, CA1, CA3 and DG for two hPSC lines (male and female n = 3 independent experiments each, data are mean ± SEM). Scale bar is 100 µm. H- Representative images of synaptic markers in hPSC-derived hippocampal neurons. Scale bar is 100 µm. I- Quantification of double-positive puncta per FOV (Field of View), SYN1 and PSD95 colocalisation in TAU+ cells for two hPSC lines (male and female, n = 3 independent experiments each, data are mean ± SEM). * *p* < 0.05, ** *p* < 0.01, \*\*\**p* < 0.001

During hippocampal development, the transcription factor BCL11B (also known as CTIP2) is expressed in postmitotic DG and CA1 neurons. Immunostaining for CTIP2 confirmed the presence of postmitotic DG and CA1-like neurons, with about 20% of CTIP2+ cells co-expressing MAP2. We also detected 12 ± 5% PROX1+ cells with about 50% of the population expressing MAP2+ confirming the presence of postmitotic DG neurons (**Supplementary Figure 3B**).

To further confirm and evaluate CA1 identity, we stained for WFS1, a marker of CA1 subfield neurons and the superficial marker, Calbindin (CALB)^59^. The CALB marker shows a subtype specific expression pattern, and it is present in glutamatergic neurons - including mature granule cells of the dentate gyrus (DG), superficial CA1 pyramidal neurons, and CA3 granule cells^60–63^. Immunostaining revealed that 50 ± 5% of cells were WFS1+, with more than 80% of this population co-expressing CALB. We next confirmed the presence of SATB2+ neurons - a marker for CA1 hippocampal pyramidal neurons which plays a role in their differentiation, by immunostaining^64,65^. Immunostaining confirmed the presence of 25 ± 2% SATB2+ cells (**Supplementary Figure 3B**).

In the human CA3 subfield, neurons are characterised by expression of ELAVL2 (HuB) and GRIK4 (KA1), which is expressed in a subset of CA3 hippocampus. At DIV 60, immunostaining confirmed 30 ± 2% ELAVL2+ cells with more than 90% of this population co-expressing MAP2. Notably, we found ELVAL2+ cells co-expressing CTIP2 (15 ± 3%) (**Supplementary Figure 3B**), indicating an immature phenotype not yet committed to a specific hippocampal subfield.

To further characterise the neuronal maturation and hippocampal subfield specification, cultures were maintained until DIV 90 with no discernible morphological difference between the male and the female cell lines. Both cell lines showed comparable cellular architecture, neurite complexity and overall culture organisation throughout the differentiation and maturation period (**Figure 2A & B**). We further characterised the cultures by immunostaining for markers specific to DG, CA1, and CA3 hippocampal subfields. Quantification focused exclusively on MAP2- and TAU+ neurons to assess subfield-specific differentiation. More than 90% of the cells expressed MAP2 and TAU (**Figure 2C** and **see Supplementary Figure 3C&D**). CTIP2 immunostaining revealed MAP2 co-expression in 50 ± 3% of CLV2-10 cell line and 30 ± 3% of CLV3-1 cell line, accompanied by a reduced proportion of PROX1+ and ZBTB20+ neurons (**Figure 2D**).

Notably, 77 ± 3% of WFS1+ cells co-expressed MAP2, confirming their neuronal identity. To further define CA1 neurons, we examined the expression of POU3F1 (OCT6), a marker enriched in the CA1 hippocampal subfield^65^. Co-expression analysis showed that 30 ± 3% of cells were WFS1+OCT6+ across both cell lines, accounting for 95 ± 3% of the total OCT6+ population (**Figure 2E**). Furthermore, more than 75% of WFS1+POU3F1+ cells co-expressed CTIP2, a transcription factor reported in CA1 neurons^66^, providing additional support for CA1 specification.Together, these findings support the specification of CA1 neurons within the differentiated cultures.

For CA3 characterisation, *GRIK4* (*KA1*), a gene enriched in CA3 neurons, was used alongside *ELAVL2*. Immunostaining revealed that 24 ± 3% of cells were ELAVL2+GRIK4+ (**Figure 2F**). Given the known molecular heterogeneity of CA3 neuronal populations and the variable expression of CA3-associated molecular markers^32,67–69^, and we noted that some of the ELAVL2+ cells did not co-express GRIK4, a gene enriched in mature CA3 neurons. This observation suggests that ELAVL2 expression alone identifies a broader CA3-associated population encompassing cells at different stages of maturation. We therefore hypothesised that a subset of these ELAVL2+GRIK4-cells may represent either immature CA3 neurons that have not yet acquired a mature molecular signature or distinct CA3 neuronal subtypes characterised by the expression of Secretagonin, *SCGN*. Immunofluorescence analyses examining the co-expression of *ELAVL2*, *SCGN*, and *GRIK4* identified both ELAVL2+SCGN+GRIK4- and ELAVL2+SCGN+GRIK4+ neuronal populations, supporting the existence of molecularly distinct CA3 subpopulations within the differentiated cultures (see **Supplementary Figure 3E**). The detection of ELAVL2+SCGN+GRIK4-cells are consistent with the interpretation that *SCGN* may mark an alternative CA3 neuronal state which do not co-express *GRIK4*^32^. Furthermore, the presence of ELAVL2+SCGN+GRIK4+ neurons corroborates our previous observations that a subset of GRIK4+ CA3 neurons also expresses *SCGN*, providing additional evidence for the existence of SCGN-expressing CA3 neuronal subtypes within the differentiated cultures. Here we demonstrate that across independent cultures with two different cells lines, the differentiation protocol reproducibly generates multiple hippocampal subfield neuronal populations comprising approximately 30-45% CA1 neurons, 25-40% CA3 neurons, and 10-20% dentate gyrus (DG) neurons (**Figure 2G**). These findings indicate that the differentiation paradigm reliably produces a heterogeneous hippocampal neuronal population containing major hippocampal subfield identities.

To assess the maturity of the cells, given that hPSC-derived hippocampal neurons are prone to slow maturation rates in vitro^70,71^, we immunostained for the presynaptic marker SYN1 and the postsynaptic marker PSD-95, observing an increased density of SYN1 puncta across both cell lines. (**Figure 2H**). We used the Synbot open-source tool^72^ to count the number of colocalised pre- and post-synaptic markers confirming the maturing state of the cultures (**Figure 2I**).

Morphologically, neurons resembling CA subfields exhibited large pyramidal somata, whereas DG-like neurons displayed smaller granule cell bodies, consistent with known hippocampal cytoarchitecture. We identified neurons which were positive for ELAVL2 and WFS1 with distinct large pyramidal somata appearing in the cultures indicating the maturation phase of the culture. Most of the neurons possessed immature dendrites with shorter processes and fewer branches (**Supplementary Figure 3C & D**).

### Generation of functional hippocampal organoids and hippocampal regionally specified astrocytes

There are reports on generating hippocampal organoids (HO) from hPSCs using embryoid bodies. The asynchronous differentiation, which leads to the presence of choroid plexus tissues, results in smaller organoids over extended culture periods, with some HO unable to survive longer than 100 days in culture^9,23^. We generated the HOs with fully committed hippocampal progenitors after confirming their identity in 2D cultures (see **Figures 1B-I**). The HOs maintained their spherical structures throughout the growth period, measuring between 750-1000 µm in diameter for both cell lines (**Figure 3A**). HOs remained free of cavities after more than 400 days in culture. Fiji-based morphometric analysis confirmed a highly uniform morphology across organoids from various cell lines, with circularity ranging from 0.945–0.992, roundness from 0.729–0.995, and solidity approaching unity (0.996–1.000) (**Figure 3A** and **Supplementary Figure 3F**). These high solidity values indicate the absence of prominent cavities or structural invaginations, consistent with the dense, uniformly dark appearance observed by brightfield imaging. Together, these data demonstrate reproducible formation of compact, non-cavitated organoids with preserved structural integrity across cultures, confirming that the differentiation protocol restricts the emergence of choroid plexus epithelial cells^23^.

**Figure 3:**
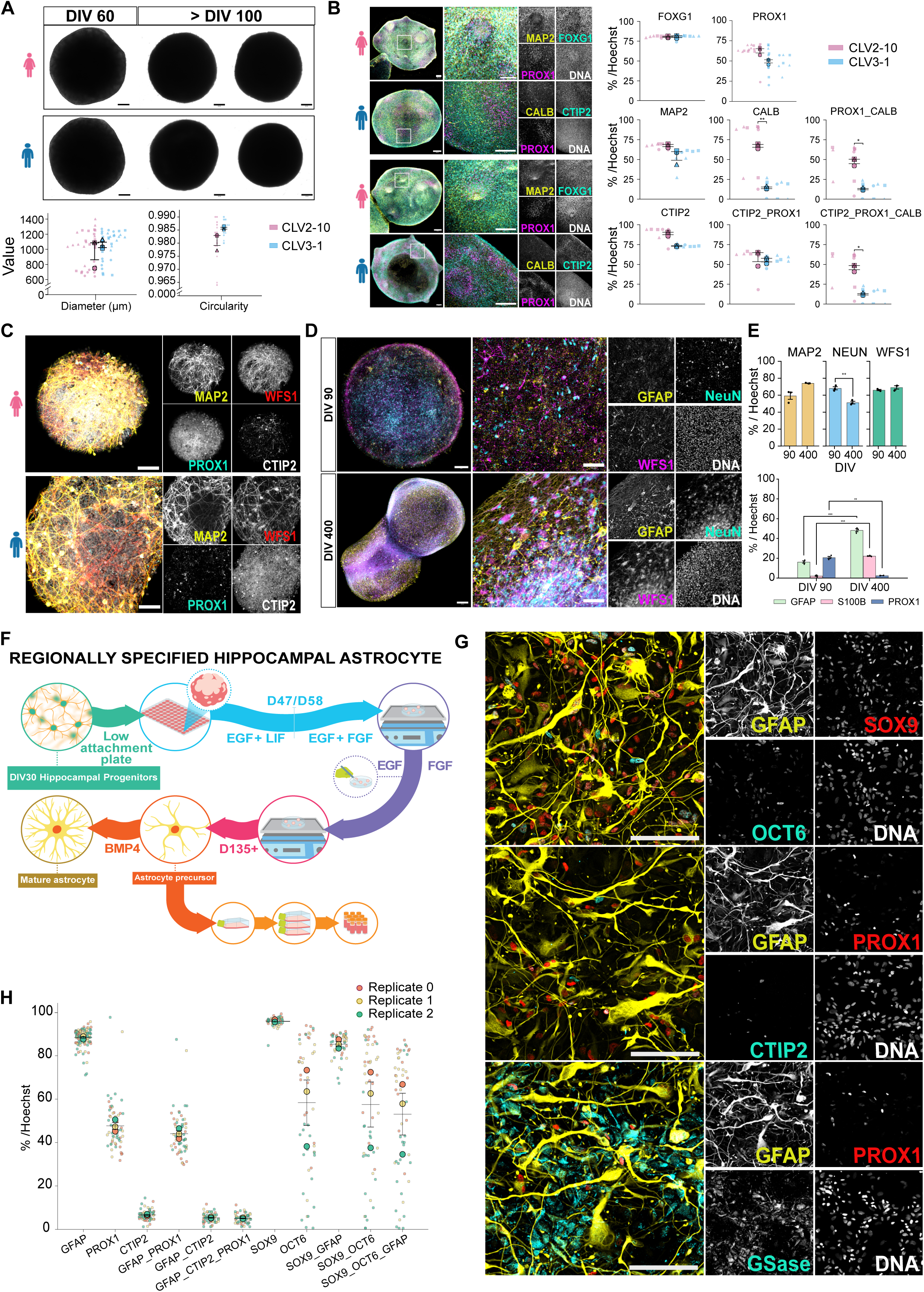
Generation of region-specific hippocampal organoids and astrocytes. A- Morphology of generated HOs at two different ages, DIV 60 and DIV 100 (left panel). Quantification of the diameter (left panel; both stages are combined) and circularity of the organoids at DIV 200 (right panel-see **Supplementary Figure 3F** for brightfield images). Scale bar is 200 µm. B- Immunostaining and quantification analysis of FOXG1+, CTIP2+, PROX1+ and MAP2+ cell population of DIV 40 HOs for two hPSC lines (male and female, n = 3 independent experiments each). The Y axis refers to the % of cells expressing the marker over the total of cells counted as per Hoechst staining. Scale bars are 200 µm. C- Immunostaining analysis of HO at DIV 200 showing CA1 and DG cell population. Scale bar is 50 µm. D- Immunostaining analysis of DIV 90 and DIV 400 HOs showing the presence of astroctyes, GFAP+ cells and NeuN+ and WFS1+ neurons. *Note that the organoids at DIV400 are two separate organoids that fused during culture.* Scale bars are E- Quantification of astrocyte-neuronal composition of HOs at DIV 90 and DIV 400 (n = 3 independent experiments, data are mean ± SEM). The Y axis refers to the % of cells expressing the marker over the total of cells counted as per Hoechst staining. F- Schematic representation of the differentiation protocol to generate hPSC-derived regionally specified hippocampal astrocytes. G- Immunostaining analysis of hPSC-derived regionally specified hippocampal astrocytes showing GFAP+, S100β+ and SOX9+ astrocytic markers, OCT6+, CTIP2+ and PROX1+ hippocampal markers and express Gluthamine Synthetase, GSase. Scale bar is 100 µm. H- Quantification of the proportion of GFAP+, S100β+ and SOX9+ astrocytic markers and OCT6+, CTIP2+ and PROX1+ hippocampal markers (n = 3 independent experiments, data are mean ± SEM). The Y axis refers to the % of cells expressing the marker over the total of cells counted as per Hoechst staining. * *p* < 0.05, ** *p* < 0.01, \*\*\**p* < 0.001

The expression of FOXG1, PROX1 and CTIP2 at DIV 40 confirmed the presence of hippocampal progenitors and DG-like neurons, with more than 50% of cells expressing MAP2 in both cell lines. CALB+ cells were present in both lines, with expression roughly 70% in the female line and 20% in the male line, consistent with the emergence of distinct neuronal subtypes within the HOs (**Figure 3B**). Immunostaining at DIV 200 confirmed the presence of PROX1+ DG neurons and WFS1+ CA1 neurons across both cell lines (**Figure 3C**). This further demonstrated the successful induction of organoids into a hippocampal fate with no significant difference between cell lines. Brain organoids can generate functional glial populations, including astrocytes and oligodendrocytes^23,73^. To assess this capacity in our HOs, we immunostained with astrocytic markers GFAP and S100β. We detected the presence of astrocytes at DIV 90 that increased progressively (GFAP, p = 0.0006; S100B, p = 0.0003; PROX1, p = 0.012; Welch’s t-test; see **Supplementary Table 12**), reaching proportions comparable to neurons by DIV 400 (**Figure 3D & E**). This further validates that the organoids can self-generate their astrocytes which are specific to the hippocampal region.

Astrocytes, distinguished by their highly ramified morphology, contribute to brain function and behaviour through homeostatic maintenance and synaptic regulation. They also play an active role in modulating neuronal network activity. These roles are mediated by their close physical proximity to neurons and dynamic interactions within neural circuits^36^. To generate regionally specified astrocytes, we followed and modified previously established protocols^74,75^. Hippocampal neural progenitor cells (NPCs) were aggregated into spheres at DIV 19 and treated with CHIR99021 from DIV 22 until DIV 35. By DIV 35, more than 30% of cells expressed the hippocampal marker PROX1, consistent with the acquisition of hippocampal identity during differentiation (refer to **Figures 1B-I**). To induce astrocytic lineage commitment, the medium was switched to astrocyte differentiation conditions, supplemented with human leukemia inhibitory factor (hLIF) and epidermal growth factor (EGF) for 4 weeks. Subsequently, cultures were maintained in medium containing EGF and fibroblast growth factor (FGF) for an additional 8-12 weeks (**Figure 3F**).

Following dissociation and plating, the cultures predominantly contained non-neuronal cells with few neuron-like morphologies. Immunocytochemical analysis revealed that the hPSC-derived astrocytes expressed canonical astrocytic markers, including glial fibrillary acidic protein (*GFAP*), S100 calcium-binding protein B (*S100*β), and *SOX9*. We also stained for gluthamine synthetase (GSase) which is highly expressed in astrocytes^76^ and the immunohistochemistry confirmed that more than 90% of the cells expressed GSase (**Figure 3G**). To assess regional identity, cells were further immunostained and quantified for hippocampal markers CTIP2, PROX1, and OCT6. A small population of CTIP2+ cells with reduced nuclear size was observed. Additionally, cells co-expressing SOX9, S100β, and GFAP also exhibited expression of OCT6, CTIP2, and PROX1, supporting the regional specification of the astrocyte population (**Figure 3G & H**).

### Temporal electrophysiological characterisation shows differences between three hippocampal culture groups

To assess neuronal functionality and network integration, three hippocampal culture groups were generated and seeded onto microelectrode arrays (MEAs): two-dimensional hippocampal neurons co-cultured with astrocytes (HN+HA), two-dimensional hippocampal neurons alone (HN), and three-dimensional hippocampal organoids (HO) (**Figure 4A**). HN+HA cultures were supplemented with astrocytes at a standard 1.4:1 neuron-to-astrocyte ratio.

**Figure 4:**
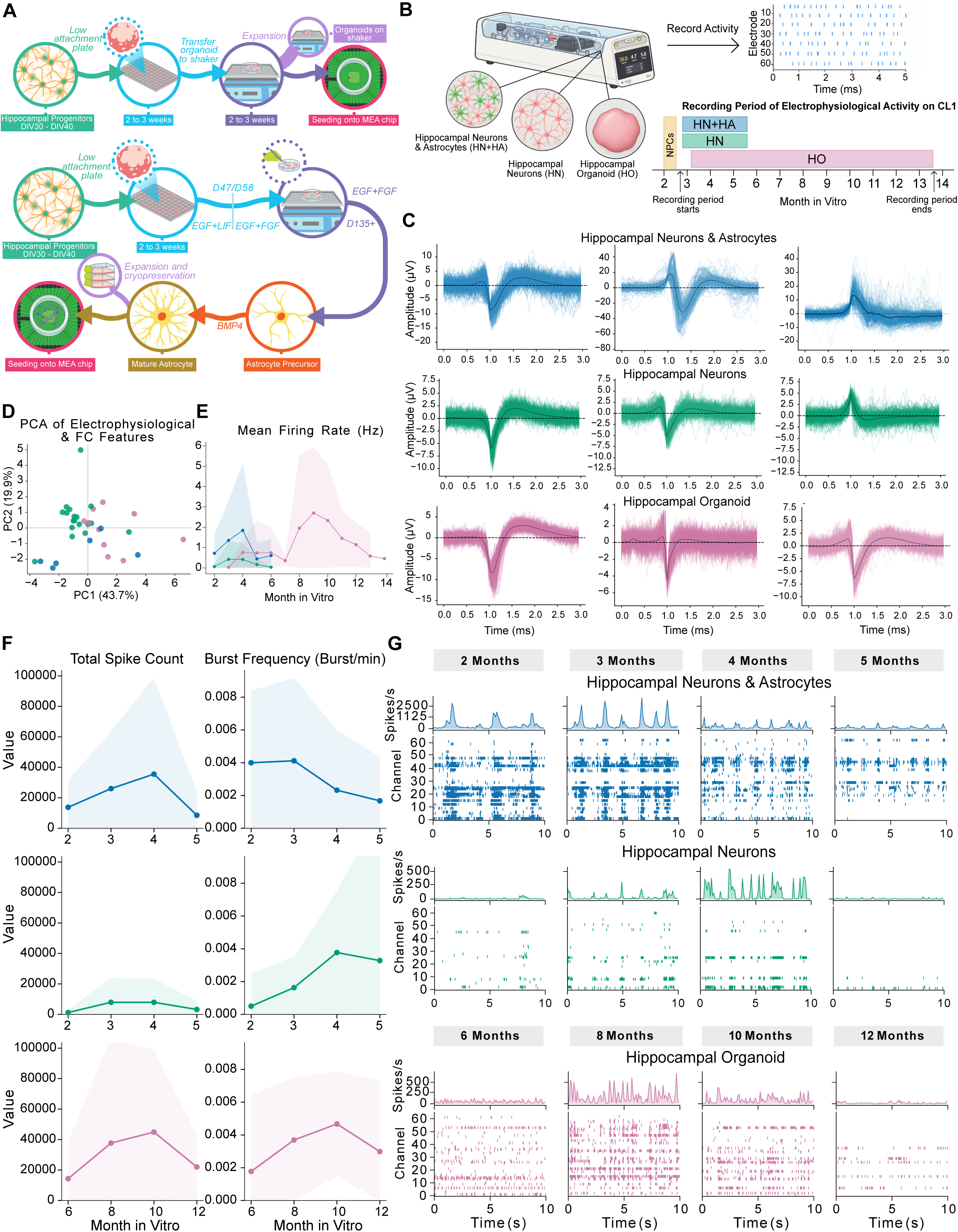
Electrophysiological characterisation of hippocampal neurons, hippocampal neuron–astrocyte co-cultures, and hippocampal organoids on the CL1 platform. A- Schematic of the differentiation and culture workflows. B- Schematic of the CL1 recording system and the three culture groups: hippocampal neurons and astrocytes (HN+HA), hippocampal neurons alone (HN), and hippocampal organoids (HO). An example raster shows spontaneous activity across electrodes. The timeline below shows the recording period for each condition in months in vitro: HN+HA and HN from approximately months 2–6 (DIV 50-160), and HO from approximately months 3–14 (DIV 60-410). C- Example waveforms of each culture group. Individual detected spikes are overlaid with the mean waveform over a 3 ms window. D- Principal component analysis of electrophysiological and functional connectivity (FC) features. Each point represents one culture, coloured by culture group type. E- Mean firing rate (Hz) across months in vitro for each culture group. Lines show the mean of all cultures per month; data are mean ± SD across cultures. F- Total spike count (left) and bursts per minute (right) over time for HN+HA (top, months 2–5), HN (middle, months 2–5) and HO (bottom, months 6–12). Lines show the mean and shaded regions indicate mean ± SD across cultures. G- Representative 10 s raster plots of spontaneous activity across MEA channels. Each is shown with its population spike-rate histogram above. Rows show HN+HA and HN at 2, 3, 4 and 5 months in vitro, and HO at 6, 8, 10 and 12 months in vitro.

Electrophysiological activity was recorded longitudinally using Cortical Labs’ CL1 system. Months in vitro (MIV) were defined from days in vitro (DIV) using 30-day intervals (i.e., DIV 0–30 = MIV 1, DIV 31–60 = MIV 2, etc.). The two-dimensional HN+HA and HN cultures were monitored from approximately 2–6 MIV (DIV 50–160), whereas the HO cultures were followed for a longer period, extending to approximately 14 MIV (DIV 410) (**Figure 4B**). Spike sorting of the recorded activity identified action-potential waveforms across all three culture groups, confirming the presence of electrically active neurons on the MEAs (**Figure 4C**).

A range of electrophysiological and Granger-causality based connectivity metrics (see **Supplementary Tables 15-16**) were measured across cultures and projected into a low-dimensional space using principal component analysis (PCA). The first two principal components captured most of the variance in the dataset (**Figure 4D**). The three culture groups showed partial separation, although some cultures from different groups exhibited similar electrophysiological profiles (**Figure 4D**). Average firing rate exhibited distinct developmental trajectories over the recording period (**Figure 4E**). All culture groups showed relatively low firing rates at the beginning of the recording period, followed by a gradual increase over the subsequent months and a decline at later timepoints. Compared with the monolayer cultures, HO cultures exhibited a delayed but more pronounced increase in firing rate, reaching a peak around 8–10 MIV before gradually declining at later timepoints. Mean firing rates were also substantially higher in HO cultures than in the monolayer groups. Among the monolayer cultures, HN+HA exhibited higher firing rates than HN, consistent with enhanced electrical activity in the presence of astrocytes^73,77,78^. The delayed timing and greater magnitude of the firing-rate increase in HO cultures are consistent with their more advanced and structurally complete functional maturation trajectory relative to two-dimensional cultures.

Total spike count and burst frequency exhibited distinct temporal trajectories across the three culture conditions (**Figure 4F**). HN+HA cultures showed a peak in total spike count around MIV 4, while burst frequency peaked around MIV 2–3, with both measures declining thereafter. Compared with HN+HA cultures, HN cultures exhibited a later increase in burst frequency, peaking around MIV 4 before subsequently declining. Despite their lower overall firing rates, HN cultures exhibited burst frequencies broadly comparable to those of HN+HA and HO cultures. HN cultures also exhibited consistently lower overall spike counts, with total spiking activity declining after approximately MIV 4. In contrast, HO cultures displayed a delayed trajectory, with total spike count peaking around MIV 8 and burst frequency reaching its peak around MIV 10, followed by a decline.

Representative raster plots and population firing-rate traces further illustrated differences in the organisation and distribution of network activity between conditions (**Figure 4G**). HN+HA cultures showed pronounced synchronisation across channels, particularly during MIV 2-3, and maintained network bursting over a longer period, whereas HN cultures exhibited comparatively sparse activity across channels and lower population firing rates. Moreover, HO cultures demonstrated widespread activity across channels and sustained population firing, with prominent clustered burst-like events evident from MIV 6–10.

### Granger-causality based connectivity analysis of dissociated hippocampal-like neurons, hippocampal neurons with astrocytes and hippocampal organoids using electrophysiology

The electrophysiological recordings revealed marked differences in spontaneous activity across the three culture groups, reflecting distinct stages of neuronal maturation and network development (**Figure 5A, Supplementary Figure 4A and B**). The HO group exhibited the highest overall neuronal excitability, with significantly higher mean firing rates and firing rate variance, together with a lower mean inter-spike interval (ISI), compared with HN cultures (mean firing rate: adjusted p = 0.008, r = 0.52; firing rate variance: adjusted p = 0.0008, r = 0.52; mean ISI: adjusted p = 0.0001, r = 0.7; see **Supplementary Table 15 and 16**). HO cultures also exhibited a higher number of network bursts and a higher burst duration ratio, indicating a greater prevalence of sustained episodes of coordinated neuronal firing rather than isolated and irregular spiking. In contrast, astrocyte-supported monolayers (HN+HA) displayed intermittent but structured firing, with moderate firing rates and reduced firing variability (**Figure 5A**). Although half of the HN+HA cultures showed no detectable network bursting, the number of network bursts among cultures bursting did not differ significantly from either HO or HN cultures, suggesting that astrocyte-supported cultures could generate recurrent network-level activity despite considerable heterogeneity in their bursting dynamics. HO cultures also exhibited a higher network burst duration ratio than the monolayer cultures, indicating that a greater proportion of their overall activity was organised into sustained network bursts. To assess the influence of extreme values, all analyses were repeated after excluding outliers, defined within each group and metric as values below Q1 − 1.5 × IQR or above Q3 + 1.5 × IQR. Outlier exclusion resulted in stronger statistical significance for several HN–HO comparisons, including mean firing rate, firing rate variance, total spike count, and maximum firing rate. (**Supplementary Figure 4D**, **Supplementary Table 17-18**), indicating that these findings were not dependent on the identified extreme values. Taken together, these findings suggest that astrocyte-supported monolayers develop organised network activity but with greater variability and less sustained bursting than HO cultures, potentially reflecting an intermediate stage of network maturation. In contrast, HN cultures exhibited minimal spontaneous activity, characterised by lower mean and maximum firing rates and lower burst frequency, consistent with less developed or weakly interconnected neuronal networks.

**Figure 5:**
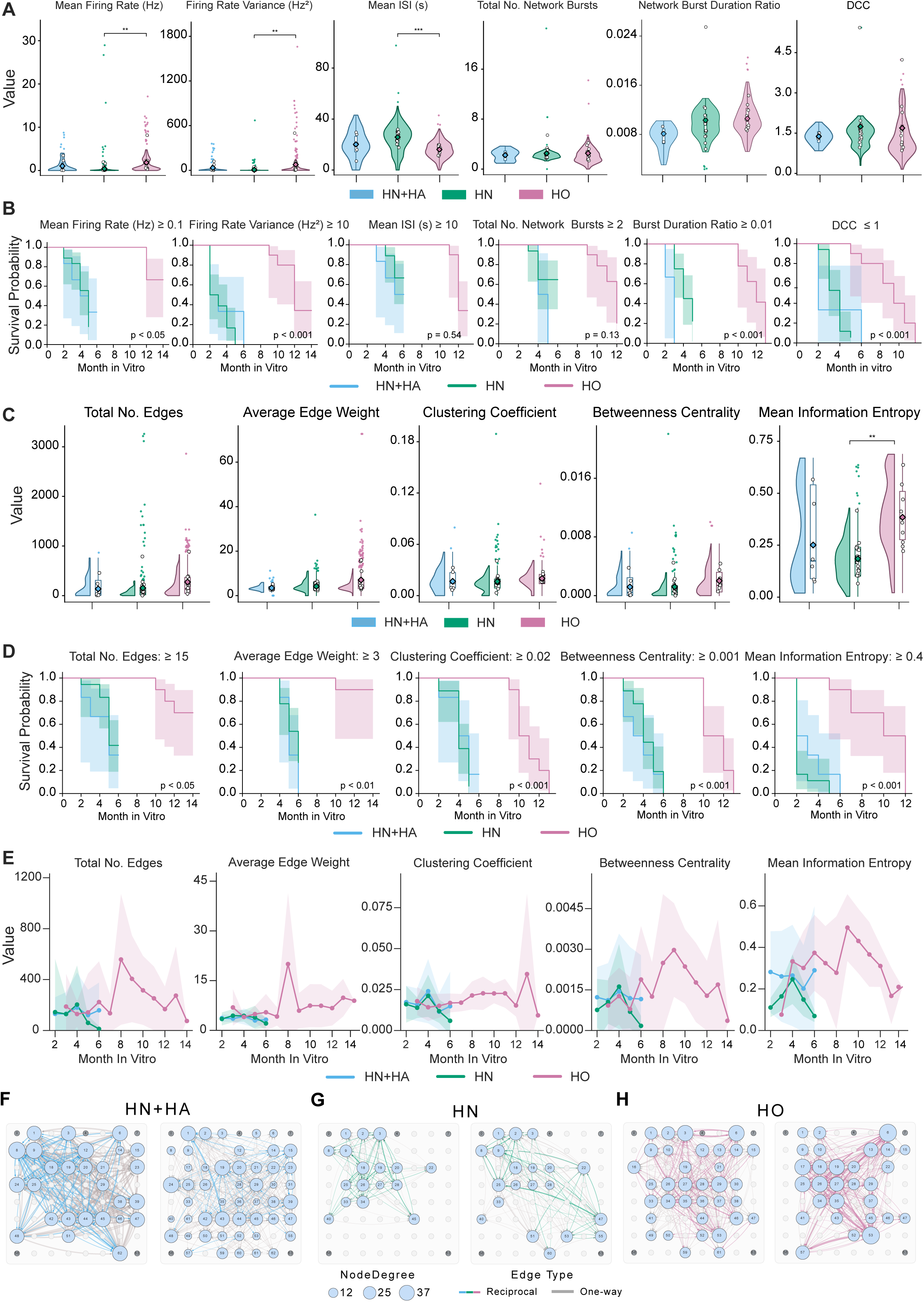
Network activity and Granger-causality based connectivity metrics differ across hippocampal culture types and change over time in vitro. A - Violin plots of electrophysiological metrics for hippocampal neurons and astrocytes (HN+HA), hippocampal neurons (HN) and hippocampal organoids (HO): mean firing rate (Hz), firing rate variance (Hz²), mean inter-spike interval (ISI, s), total number of network bursts, network burst duration ratio, and deviation from criticality coefficient (DCC). Each violin shows the distribution of values across recordings, with culture-level means indicated. Statistical comparisons between groups were performed on culture-level means using the Kruskal–Wallis test followed by Dunn’s post hoc test with Bonferroni correction; *p < 0.05, **p < 0.01, ***p < 0.001 (see **Supplementary Tables 14 & 15**). B- Kaplan–Meier survival curves showing culture longevity as a function of meeting predefined functional activity thresholds for each metric shown in (A). Survival probability is plotted against days in vitro (DIV) for each culture group. Thresholds were chosen to reflect minimal functional or network-level activity: mean firing rate ≥ 0.1 Hz, firing rate variance ≥ 10 Hz², mean ISI ≥ 10 s, burst duration ratio ≥ 0.01, total network bursts ≥ 2, DCC ≤ 1. Cultures failing to meet the specified threshold were considered to have failed the criterion. Shaded regions indicate 95% confidence intervals. Across nearly all metrics, HO cultures exhibit significantly prolonged survival and sustained functional activity compared to HN or HN+HA cultures. C - Raincloud plots of Granger-causality based network metrics and mean information entropy for hippocampal culture groups: total edge counts, average edge weights, clustering coefficient, betweenness centrality and mean information entropy. Each raincloud shows the distribution of values across recordings, with culture-level means indicated. Statistical comparisons between groups were performed on culture-level means using the Kruskal–Wallis test followed by Dunn’s post hoc test with Bonferroni correction]; *p < 0.05, **p < 0.01, ***p < 0.001 D - Kaplan–Meier survival curves showing culture longevity as a function of meeting predefined functional activity thresholds for each metric shown in (C). Survival probability is plotted against days in vitro (DIV) for each culture group. Thresholds: total edge counts ≥ 15, average edge weight ≥ 3, clustering coefficient ≥ 0.02, betweenness centrality ≥ 0.001, and mean information entropy ≥ 0.4. Cultures failing to meet the specified threshold were considered to have failed the criterion. Shaded regions indicate 95% confidence intervals. E - Changes over time in total edge count, average edge weight, clustering coefficient, betweenness centrality and mean information entropy from months 2–14 in vitro. Lines and points show the mean across cultures per month, and shading shows ± 1 SD. F-H - Representative Granger-causality based network graphs for HN+HA (F), HN (G) and HO (H). Nodes represent individual electrodes on the 64-channel MEA, labelled by channel number. Blue nodes represent active electrodes, dark grey nodes represent reference or ground electrodes. Node size shows node degree. For clarity, only the top 300 edges (ranked by edge weight) are shown. Edges are either reciprocal or one-way connections.

To further assess network stability and information-processing dynamics, we quantified the deviation from criticality coefficient (DCC) for each culture (see **Supplementary Methods** for details). (**Figure 5A**). Previous work has shown that exposure to structured, task-positive environments can shift in vitro neuronal networks toward dynamical states closer to criticality, where information transmission and adaptive behaviour may be enhanced^79^. In contrast, under the spontaneous recording conditions used in this study, population dynamics across all groups remained displaced from the critical regime, potentially reflecting the absence of externally applied stimulation. Despite this overall deviation from criticality, HN+HA cultures exhibited the lowest mean DCC, indicating a descriptive tendency toward dynamics closer to the critical regime than HN and HO cultures, although this difference did not reach statistical significance. In contrast, HO cultures showed an intermediate mean DCC with a broader distribution, suggesting greater heterogeneity in their network dynamics.

Survival analyses further demonstrated that the HO group maintained functional electrophysiological thresholds across multiple metrics for significantly longer relative time periods compared to the other groups (log-rank tests; p < 0.05 for mean firing rate, and p < 0.001 for firing rate variance, burst duration ratio, and total network bursts, see **Supplementary Table 19-20**), while monolayer cultures rapidly dropped below these thresholds, with no significant difference between the two monolayer groups (**Figure 5B and Supplementary Figure 4C**). Together, these results demonstrate progressive functional maturation across the three models, with organoid cultures showing the most robust and integrated electrophysiological signatures, astrocyte-supported monolayers displaying intermediate functionality, and pure neuronal monolayers exhibited more limited electrophysiological development.

### Granger-causality based connectivity and network topology

Next, network dynamics and directed functional interactions among electrodes were examined using complementary measures of information and Granger-causality based connectivity. Mean information entropy was quantified to assess the variability and complexity of neuronal activity across electrodes, providing a measure of the information content within spontaneous network dynamics. Directed functional interactions were subsequently examined using Granger causality–based connectivity graphs (see **Supplementary Methods** for details). This approach quantifies the extent to which the activity of one electrode can predict that of another, allowing inference of directed, weighted network architectures underlying spontaneous activity.

Granger causality–derived networks differed qualitatively across culture conditions in topology, connectivity strength, and temporal stability (**Figure 5C**). However, aggregated network statistics showed no significant differences between culture groups for any metric examined except mean information entropy, which was significantly higher in HO than in HN cultures (adjusted p = 0.002, r = 0.58; **Supplementary Tables 21-22**). This suggests greater complexity and diversity of spontaneous activity in HO cultures.

Because the HN group contained a relatively large number of potential outliers, a sensitivity analysis was performed, excluding values beyond 1.5 × IQR. After exclusion of outliers, the difference in mean information entropy became more pronounced, and HO cultures also showed significantly higher betweenness centrality (adjusted p = 0.019, r = 0.47), total number of edges (adjusted p = 0.011, r = 0.50), and clustering coefficient (adjusted p = 0.03, r = 0.43) than HN cultures (**Supplementary Figure 4E; Supplementary Tables 23-24**). These results point to greater overall connectivity, local clustering, and centralisation of network interactions in HO cultures. Because these differences emerged only after outlier exclusion, they should be interpreted with caution, although they suggest that HN and HO cultures may differ in aspects of network organisation, with HO networks exhibiting greater overall connectivity, local network clustering and centralisation of network interactions. Although not statistically significant, HN+HA cultures appeared to occupy an intermediate regime, with network metrics generally falling between those of HN and HO cultures.

To assess the temporal persistence of network organisation, we performed survival analyses based on thresholded network metrics (**Figure 5D**). HO cultures maintained high levels of connectivity (i.e. total edge counts), average edge strength, clustering, mean information entropy and betweenness centrality for significantly longer relative time periods compared to both monolayer conditions (log-rank tests; p-values correspond to total edge count, average edge weight, clustering coefficient, betweenness centrality, and mean information entropy, respectively: HO vs HN, total edge count p < 0.05, average edge weight p < 0.01, and clustering coefficient, betweenness centrality, and mean information entropy all p < 0.001; see **Supplementary Table 25-26**), indicating sustained preservation of complex network structure. Additionally, across some metrics, HN+HA cultures exhibited intermediate stability, more closely resembling HN cultures, although these differences were not statistically significant, suggesting a graded pattern of network organisation across the three culture conditions.

Finally, analysis of connectivity metrics aggregated into monthly bins across days in vitro revealed distinct developmental trajectories in network topology (**Figure 5E**). Organoid networks exhibited prolonged and dynamic reconfiguration, with gradual increases and sustained variability in edge counts, clustering, centrality and information entropy measures over extended developmental windows. Monolayer cultures, by comparison, showed earlier stabilisation of network properties, with HN+HA cultures maintaining greater total edge counts, clustering coefficient, betweenness centrality, and mean information entropy. These patterns were also apparent in the representative Granger causality–derived connectivity network graphs, where HN+HA and HO cultures exhibited greater numbers of edges and higher node degrees, together with stronger edge weights, compared with HN cultures (**Figure 5F-H**).

Together, these results demonstrate that three-dimensional organoid architectures support the emergence, persistence, and continued reorganisation of complex directed causal and functional networks, whereas two-dimensional monolayers converge more rapidly toward simpler and more stable connectivity regimes.

### Bulk transcriptomics reveals a reproducible developmental trajectory

We used bulk transcriptome analysis to analyse the transcriptional changes associated with hippocampal differentiation and maturation across six developmental stages spanning DIV 0 to DIV 200 (see **Supplementary Table 27**). Gene-level read counts were compiled into a count matrix comprising 17 bulk RNA-seq samples and 26,293 genes. Principal-component analysis (PCA) of the bulk RNA-seq data revealed a clear temporal progression of transcriptomic states, with all differentiation stages separating predominantly on the PC1 axis and following a continuous developmental trajectory from neural progenitor cells (DIV 0-11) to mature neuronal cultures (DIV 154-200). Samples were arranged along a continuous trajectory according to culture stage, progressing from the NPC stages (DIV 0-11) to the latest organoid stages (DIV 154–200), with replicates from the same stage grouped together (**Figure 6A**). In line with previous bulk transcriptomic studies, classical housekeeping genes (GAPDH, ACTB, UBC, HSPD1, TUBB, HDAC1, TBP, and VDAC1) exhibited only modest expression changes relative to DIV 0 throughout the differentiation time course, highlighting their limited suitability as reference genes for studies investigating dynamic developmental processes (**Figure 6B**). In contrast, expression of Cannabinoid receptor 1 (CNR1), increased progressively across successive culture stages, following the developmental trajectory observed in the global transcriptomic analyses. The progressive enrichment of CNR1 across differentiation stages provides independent molecular validation of ongoing neuronal maturation and developmental progression in vitro^80^ (**Figure 6C**).

**Figure 6:**
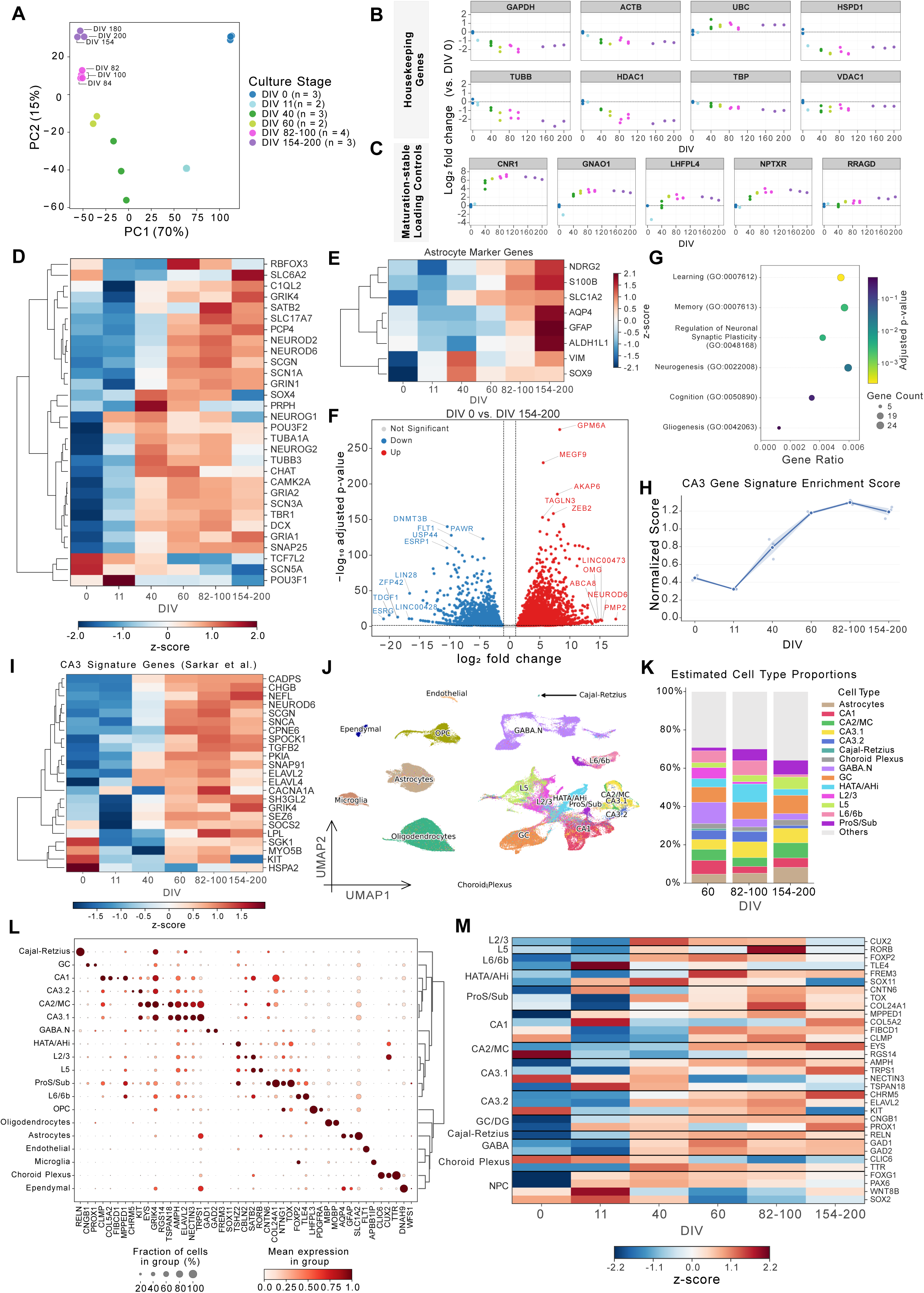
Bulk transcriptomic signatures align with hippocampal snRNA-seq references. A - PCA of bulk RNA-seq profiles at DIV 0 (n = 3), DIV 11 (n = 2), DIV 40 (n = 3), DIV 60 (n = 2), DIV 82–100 (n = 4) and DIV 154–200 (n = 3) B - Log₂ fold change (relative to DIV 0) in expression of the housekeeping genes GAPDH, ACTB, UBC, HSPD1, TUBB, HDAC1, TBP and VDAC1 across DIV. Each point is one sample, coloured by culture stage. The dashed line indicates no change. C - Log₂ fold change (relative to DIV 0) in expression of the maturation-stable loading controls CNR1, GNAO1, LHFPL4, NPTXR and RRAGD across DIV. D - Heatmap of z-scored expression of neuronal and neurodevelopmental marker genes across culture stages. Genes are hierarchically clustered. E - Heatmap of z-scored expression of astrocyte marker genes (NDRG2, S100B, SLC1A2, AQP4, GFAP, ALDH1L1, VIM and SOX9) across culture stages. Genes are hierarchically clustered. F - Volcano plot of differential expression between DIV 0 and DIV 154–200. Genes significantly upregulated at DIV 154–200 are shown in red, significantly downregulated genes in blue, and non-significant genes in grey (adjusted-p < 0.05, |log₂ fold change| > 1). For each direction, the top 5 significant genes by adjusted p-value and the top 5 by log₂ fold change are labelled. G - Gene Ontology (GO) enrichment of genes that are upregulated at DIV 154–200, highlighting biological processes associated with hippocampal function. The x-axis shows the gene ratio, point size shows gene count, and colour shows adjusted p-value. H- Normalized CA3 signature enrichment scores across culture stages, calculated as the mean z-score of CA3 signature genes. The line shows the mean, the shaded band shows ± SEM, and points show individual samples. The dashed line indicates a score of zero. I- Heatmap of z-scored expression of 23 CA3 signature genes across culture stages. J- UMAP of the reference single-nucleus RNA-seq dataset of the human hippocampus used for deconvolution, coloured by annotated cell type. K- Estimated cell-type proportions at DIV 60, DIV 82–100 and DIV 154–200, obtained by deconvolution of bulk RNA-seq data using BisqueRNA with the snRNA-seq reference in (J). L- Dot plot of the expression of representative marker genes across cell types in the snRNA-seq reference in (J). Marker genes shown are RELN (Cajal–Retzius); CNGB1 and PROX1 (GC); CLMP, COL5A2, FIBCD1 and MPPED1 (CA1); EYS and RGS14 (CA2/MC); KIT, GRIK4, TSPAN18, AMPH, NECTIN3, TRPS1, CHRM5 and ELAVL2 (CA3.1 and CA3.2); GAD1 and GAD2 (GABA.N); FREM3 and SOX11 (HATA/AHi); CBLN2 and CUX2 (L2/3); SATB2 (CA1 and L2/3); TSHZ2 (ProS/Sub, L2/3 and HATA/AHi); RORB (L5); FOXP2 and TLE4 (L6/6b); CNTN6, COL24A1, NTNG1, TOX (ProS/Sub); LHFPL3 and PDGFRA (OPC); MBP and MOBP (oligodendrocytes); AQP4, GFAP and SLC1A2 (astrocytes); FLT1 (endothelial); APBB1IP (microglia); CLIC6 and TTR (choroid plexus); and DNAH9 (ependymal). SATB2 marks both CA1 and cortical neurons, as it is expressed in CA1 pyramidal neurons and upper-layer cortical projection neurons. SATB2 marks both CA1 and cortical neurons, as it is expressed in CA1 pyramidal neurons and upper-layer cortical projection neurons. Its absence from the HATA/AHi, CA2/MC and CA3 clusters supports the separation of CA1 and the cortical subpopulations from these cell types. Dot size indicates the fraction of cells expressing each gene, while colour indicates scaled mean expression. Cell types are hierarchically clustered. M-Heatmap of z-scored expression of selected representative marker genes for each cell type across culture stages in the bulk RNA-seq data, grouped by cell type. FOXG1, PAX6, WNT8B and SOX2 were used as markers of neural progenitor cells (NPCs).

Marker gene analysis revealed a transcriptional progression from early neurogenesis to late-stage neuronal and glial maturation during hippocampal differentiation **(Figure 6D)**. Early neurogenic markers, including NEUROG1, NEUROG2, SOX4, TUBB3, and DCX, were enriched during the initial stages of differentiation before declining at later time points, while POU3F1 peaked at DIV 11, consistent with its role in early neuronal specification. Markers associated with neuronal maturation and excitatory neuronal identity, including SATB2, SCN1A, and SNAP25, increased progressively and remained highly expressed from DIV 60 onwards, indicative of continued neuronal maturation.

Consistent with the developmental transition from neurogenesis to astrogliogenesis during culture maturation, expression of astrocytic markers emerged later during the differentiation becoming markedly enriched at DIV 82-100 and DIV 154-200 (**Figure 6E**) validating the immunostaining analysis of the organoids at DIV 90 (see **Figure 3D & E**). Consistent with these observations, differential expression analysis comparing DIV 0 and DIV 154-200 demonstrated significant enrichment of genes associated with neuronal maturation, membrane organisation, synaptic signalling, and glial development at later stages, whereas genes associated with pluripotency and early developmental states were significantly downregulated (adjusted p < 0.05; **Figure 6F, see Supplementary Table 36**). Collectively, these findings indicate that the differentiation paradigm recapitulates key transcriptional features of hippocampal development and supports the progressive acquisition of mature neuronal and glial identities in vitro.

To determine whether transcriptional maturation was accompanied by the acquisition of hippocampal functional signatures, gene ontology (GO) enrichment analysis was performed on developmentally regulated genes. Enriched terms included learning, memory, cognition, and astrocyte differentiation, all of which are associated with hippocampal development and function (**Figure 6G**, **Supplementary Figure 5B** and **Supplementary Tables 47-51**)^81^. Together, these findings demonstrate the progressive acquisition of transcriptional signatures associated with mature hippocampal neuronal and glial identities, consistent with the emergence of functional hippocampal maturation during long-term differentiation.

To further assess CA3 regional specification, we correlated the bulk RNA-seq profiles with a published CA3 pyramidal neuron gene signature^32^. CA3 signature scores increased markedly after DIV 40 and plateaued from DIV 82-100 onwards, consistent with the progressive establishment and maintenance of a stable CA3-like transcriptional identity (ELAVL2, SCGN, GRIK4) during long-term differentiation (**Figures 6H-I**).

To estimate cellular composition, bulk RNA-seq profiles were deconvolved using a high-resolution human hippocampal single-nucleus reference atlas integrating postnatal and adult datasets (**Figures 6J**, **Supplementary Figure 5 and 6**)^82^. To resolve the glutamatergic neuronal cluster at finer resolution, cells within the glutamatergic cluster were re-annotated by label transfer from the adult human hippocampal snRNA-seq dataset containing detailed glutamatergic subtype labels^65^. Deconvolution revealed a progressive increase in hippocampal excitatory neuronal populations, including CA-field and dentate gyrus neurons, accompanied by a decline in Cajal-Retzius cells, consistent with developmental maturation (**Supplementary Tables 52-54**)^83^. These trends were supported by the temporal enrichment of region-specific hippocampal markers, ProS/Sub (CNTN6, TOX, COL24A1), CA1 (MPPED1, COL5A2, FIBCD1), CA3 (AMPH, TRPS1, CHRM5) and DG (PROX1, CNGB1)^32,84^ and the concomitant reduction of progenitor-associated genes (FOXG1, PAX6, WNT8B, SOX2)^32^, indicating the progressive emergence of mature and regionally specified hippocampal identities in vitro (**Figures 6K-M** and **Supplementary Figure 5, 6 & 7**).

### MPPED1 reporter validates CA1 neuron identity

To functionally validate the generation of CA1 neurons, a population that remains challenging to robustly identify in existing hippocampal differentiation protocols^9,19,25,26^, we developed an MPPED1 promoter-driven fluorescent reporter strategy (**Figure 7A**). The gene, *MPPED1* was selected based on its progressive enrichment during differentiation and its association with CA1 identity across transcriptomic datasets^83,84^ (**Figure 7B & C**). To maximise signal specificity and facilitate quantitative analysis, the reporter construct was designed to express a histone H2B-tagged TdTomato fluorescent protein (H2B-TdTomato), thereby restricting reporter localisation to the nucleus. Nuclear confinement produces a discrete and readily quantifiable fluorescent signal while improving compatibility with fixation, immunohistochemistry, and cell-counting analyses^65,82^ (**Figure 7A**). Reporter activation was readily detected following lentiviral transduction, and quantitative co-localisation analyses demonstrated overlap with the canonical CA1 markers WFS1 and OCT6 (**Figure 7D**). Together, these findings provide functional validation of CA1 neuron generation and further support the ability of this differentiation protocol to produce all major hippocampal subfield neuronal populations.

**Figure 7:**
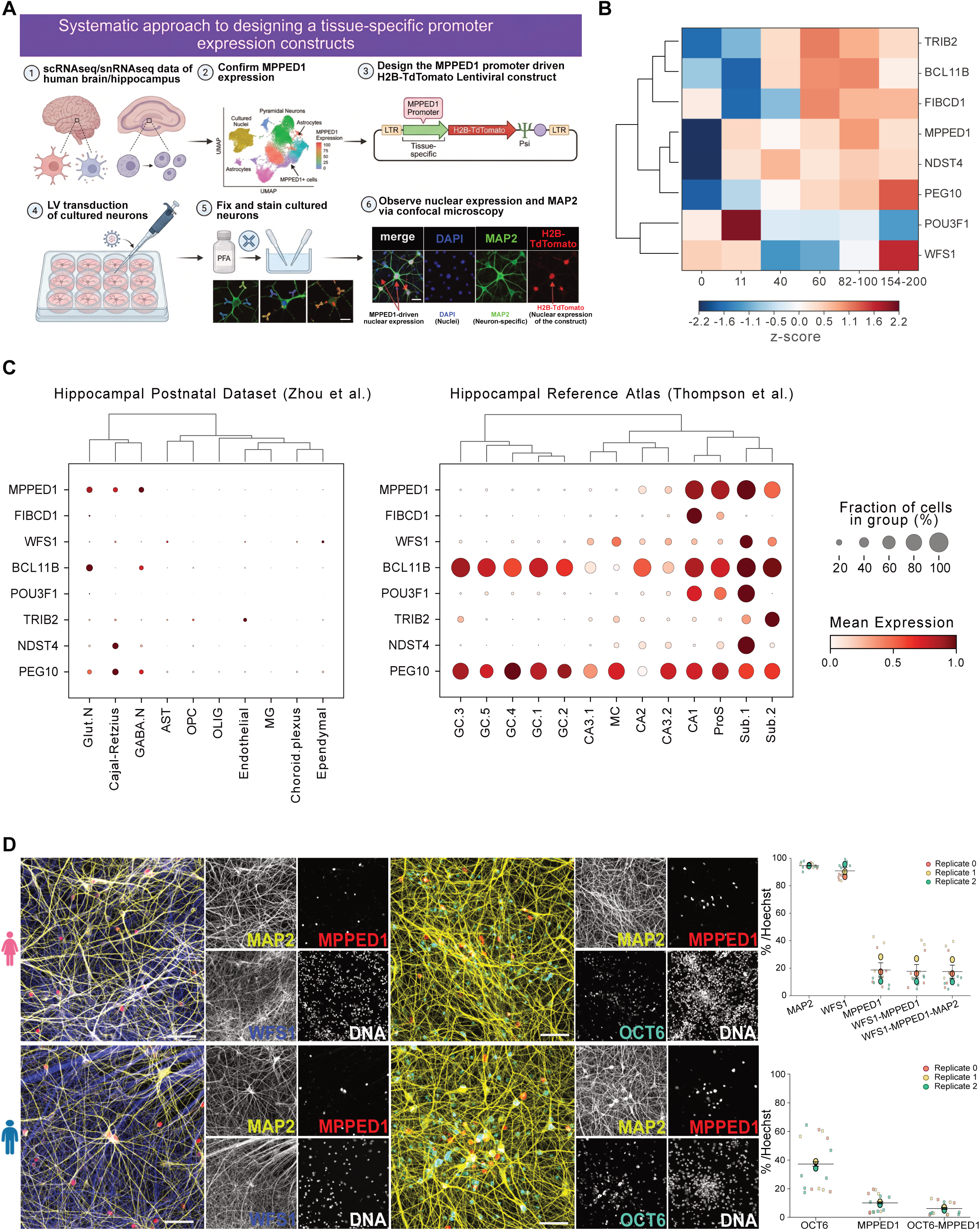
MPPED1 reporter functionally validates CA1 neuron specification. A- Graphic representation of the approach to design MPPED1 hippocampal subfield population expressing construct. B- Heatmap showing z-scored expression of genes enriched in the CA1 neuronal population. Marker selection was based on enriched CA1 genes previously identified. Genes are hierarchically clustered based on expression patterns. C- Dot plot illustrating the expression of representative CA1 marker genes across cell types in a human hippocampal snRNA-seq reference dataset. D- Immunostaining and quantitative analysis of long-term cultures transduced with the MPPED1-H2B-TdTomato reporter demonstrating co-localisation of reporter-positive cells with the CA1 markers POU3F1 (OCT6) and WFS1 across two hPSC lines (male and female n = 3 independent experiments each, data are mean ± SEM). Scale bar is 100 µm.

## Discussion

In this study, we have demonstrated a robust and modular approach for the generation of dorsomedial telencephalic region progenitors from hPSCs through a modified tri-inhibitory induction method^46,85^. Introducing the FGF-MAPK signalling by treating the cells with FGF2 led to the maintenance of the progenitor phenotype and the expression of TBR2+ IPCs, which play a role in hippocampal progenitor fate specification. This study established a protocol which demonstrates the increased expression of PROX1+ cells in cultures. When cultured over time, these progenitors gave rise to all key hippocampal subtypes, including dentate gyrus (DG), CA3 neurons, and – until now – the elusive CA1 neuron subtypes. While earlier protocols enabled DG and CA3 neural differentiation, until this work, a reliable protocol that generates a relatively meaningful number of neurons with a CA1 identity has been notably absent^9,19,25,26^. Given the known critical role of the CA1 neural subtype in integrating and outputting processed information, this has been a significant limitation in understanding the dynamics of hippocampal cells that can now be overcome^5,6,12,13^. Moreover, it was established that the range of subtypes is not a feature of the donor line, as the results were replicated with two independent hPSC lines from different donors. It is proposed that future research will be able to use the protocol described in this work to explore and better understand the computational properties of hippocampal neural cultures with a range of subtypes^86,87^.

Mechanistically, the staged manipulation of signalling pathways appears central for the protocol to guide early dorsomedial identity toward hippocampal lineages. Early tri-inhibition, combined with SHH pathway inhibition to prevent ventralisation^48,49^, produced high proportions of dorsal forebrain progenitors, consistent with a dorsomedial telencephalic starting point. Subsequent exposure to FGF2 was associated with maintenance of a progenitor phenotype and emergence of TBR2+ intermediate progenitor cells, a lineage state implicated in hippocampal neurogenesis and the progression toward PROX1+ DG programs in vivo. This intermediate progenitor phase likely creates a permissive developmental window in which WNT activation (via CHIR99021) can drive hippocampal progenitor programs, including sustained increases in PROX1 expression over time^33,34^. Notably, the observed sensitivity to CHIR99021 dose is consistent with WNT’s dual role in regulating proliferation and fate decisions^30^. These results emphasise that controlled WNT activation can be used in a tuneable manner, as a morphogen that shapes progenitor expansion and downstream differentiation trajectories.

Along with standard monolayers, organoids were also generated, as were hippocampal regionally specified astrocytes to allow the astrocyte proportion of the cultures to be further enhanced to more physiologically relevant levels of expression. These neural cultures survived and were electrically active for extended periods of time, in the case of the organoids, over 400 days. This supports the idea that these are a useful model for long-term assessment for neurodevelopment, toxicology and pharmacology, and for studying neural dynamics. These cultures were electrically active networks, with functional synaptic connections identified, as confirmed by electrophysiological activity and immunocytochemical analysis. Of particular interest was the long-term and detailed comparison of nuanced electrophysiological activity across the different culture types. The firing rate data demonstrated that organoids persisted in electrophysiological activity for longer and exhibited significantly higher mean firing rates than monolayers. However, peak firing rates did not follow this pattern, in some instances, hippocampal neuron–astrocyte monolayers exhibited peak firing rates comparable to or even higher than those of organoids. Organoids also initially displayed dynamics closer to criticality than the monolayer cultures, although this advantage diminished over time as their dynamics drifted further from the critical state. This is consistent with recent work showing that organoids can sustain near-critical dynamics more effectively than monolayers, but that structured stimulation is required for a culture to remain in a critical state^79,90^. Another notable finding was that mean information entropy was significantly higher in hippocampal organoids than in hippocampal monolayers. Additionally, sensitivity analysis following the exclusion of extreme values revealed significantly higher values for almost all Granger-causality-derived network metrics in hippocampal organoids compared with hippocampal monolayers, with particularly pronounced differences in betweenness centrality and total edge counts.

Hippocampal neuron–astrocyte cultures showed intermediate values across these network metrics, and although these did not differ significantly from either hippocampal monolayers or organoids, the consistent trend towards higher connectivity than monolayers suggests that astrocyte co-cultures partially enhance network integration, with the lack of significance likely reflecting the high variability observed within this group. Overall, while the addition of astrocytes appeared to partially enhance electrophysiological activity and network connectivity in monolayers, these results suggest that full maturation and network integration are better achieved in the organoid model than in traditional monolayers.

It is also noteworthy that region-specific astrocytes could be reliably generated with only minor modifications to the original protocol. A growing body of literature has identified that astrocyte function is region-specific, with key differences in how these glial cells interact and support neurons^35–38^. For future work that aims to understand the functional interactions between astrocytes and neurons, along with how different neuron-astrocyte ratios may influence network dynamics, the ability to more tightly control the proportions of these cells in a region-specific manner will be valuable. A detailed analysis of these aspects was beyond the scope of the present study, which focused on differentiation methods and their validation. Nevertheless, several electrophysiological differences between standard monolayers and those supplemented with additional astrocytes were apparent. Astrocyte-supplemented cultures displayed a higher mean firing rate, shorter bursts, and increased edge weights and betweenness centrality. Uncovering the mechanisms underlying these functional differences will be an important direction for future research.

By leveraging snRNAseq dataset and our bulk RNAseq dataset to generate a MPPED1 promoter-driven H2B-TdTomato reporter, we developed a MPPED1 promoter-driven H2B-TdTomato reporter that enables the selective identification and quantification of CA1 neurons. The progressive emergence of MPPED1 during differentiation, together with the strong colocalisation of reporter-positive cells with the canonical CA1 markers WFS1 and POU3F1, provides independent validation of CA1 identity. Together with our molecular and phenotypic analyses, these findings demonstrate that the differentiation paradigm reproducibly generates the three major hippocampal subfield populations, including CA1, CA3, and DG neurons. More broadly, the development of a CA1-specific reporter system provides a valuable tool for the prospective identification, isolation, and functional interrogation of human CA1 neurons, thereby expanding the utility of this platform for studies of hippocampal development, circuit formation, and disease modelling.

A key limitation of this work however is that in terms of cell culture structure, it only compared organoids to monolayers. As such, this work cannot conclusively answer whether these electrophysiological properties are a result of the organoids having a specific spherical morphology, having greater cell density, or some other feature. Future work will be well suited to explore in greater detail the structure-function relationship with these cell types, including the use of modular cell types that specifically control geometry and topology of these cultures^91^. Another limitation is that although marker-based immunostaining supports the presence of DG, CA1, and CA3 neurons, the observed overlap in some marker combinations and the persistence of immature dendritic morphology indicate that a subset of cells may remain in transitional states. It is also possible that the cellular identity may not yet be fully resolved for all neurons even at later timepoints. Secondly, while MEA recordings demonstrate network activity and maturation differences across culture formats, they do not directly resolve synapse type (excitatory versus inhibitory), cell-type-specific contributions, or trisynaptic-like circuit motifs. Incorporating targeted interneuron differentiation, more explicit excitatory/inhibitory balancing, and stimulation paradigms (including patterned theta-like stimulation) would strengthen links between the observed network behaviours and how the hippocampal functions are central to learning and memory.

Ultimately, this work has demonstrated an improved protocol for generating all major hippocampal subtypes from hPSCs. Not only does this approach generate the long-elusive CA1 subtype that has been missing from previous protocols, but the method is able to produce region-specific astrocytes. The rich electrophysiological properties of the neural cultures were found to be sensitive to morphology (monolayer vs organoid) and astrocyte to neuron ratio. The hippocampus, located in the medial temporal lobe, engages in bidirectional communication with the cerebral cortex, and these interactions are essential for memory formation, consolidation, and retrieval^90^.

Future studies integrating hippocampal and cortical populations will provide an opportunity to investigate the bidirectional interactions which is a prerequisite for hippocampal-cortical communication in vivo. Although much remains to be understood regarding hippocampal cell-type diversity and inter-regional circuitry, the ability to generate and maintain diverse neural populations in vitro provide a valuable platform for dissecting the cellular and functional mechanisms governing human brain development, connectivity, and disease.

## Methods

### hPSC maintenance and differentiation

hPSC lines CLV2.10 (CLV2-10), CLV3.1 (CLV3-1) and RM3.5 TdTomato, were maintained as previously described^91^. Hippocampal induction was achieved using an established cortical induction protocol with adaptations^46,94^. hPSCs were seeded into Laminin-521-coated (5 μg/mL) plates at 0.15 × 10^6^/cm^2^ in mTeSR1 medium with 50 nM Chroman 1 (MedChem Express). Twenty-four hours later, cells were switched to dual-SMAD and Wnt inhibition for 6 days in basal differentiation medium (hereby referred to as CTX medium) supplemented with 100 nM LDN193189 (Stemgent), 10 μM SB431542 (R&D Systems) and 2 µM XAV-939 (MedChemExpress). The CTX medium consisted of 1:1 DMEM/F12 and Neurobasal with 0.5 × B27, 0.5 × N2, 0.5 × ITSA, 0.5 × GlutaMAX, 0.5 × penicillin/streptomycin, and 50 μM 2-mercaptoethanol (Life Technologies). On DIV 2, media was additionally supplemented with a hedgehog inhibitor (3 µM Cyclopamine; MedChemExpress) to inhibit ventralisation of the progrenitors. On DIV 7, Wnt inhibitor XAV-939 is removed. On DIV 11, progenitors were passaged with 0.5 µM EDTA in DPBS (without calcium and magnesium) and reseeded at 1:2.5x supplemented with Chroman 1. The following day, cultures were transferred to CTX containing 20 ng/mL fibroblast growth factor 2 (FGF2; R&D Systems) and 3 µM Cyclopamine for 8 days. On DIV 19, progenitors were passaged with accutase and counted using the haemocytometer. Progenitors were cryopreserved or further differentiated into hippocampal progenitors and neurons.

To differentiate cortical progenitors, we followed the Gantner *et al.* protocol to achieve DIV 19 progenitors^45^. Briefly, cells were differentiated for 11 days using CTX media supplemented with 100 nM LDN193189 and 10 μM SB431542. On DIV 11, progenitors were passaged and reseeded at 1:2.5x supplemented with Chroman 1. The following day, cultures were transferred to CTX containing 20 ng/mL fibroblast growth factor 2 (FGF2; R&D Systems) for 8 days. On DIV 19, progenitors were passaged and counted using the haemocytometer. Progenitors were cryopreserved or further differentiated into cortical progenitors and neurons.

### Differentiation into hippocampal progenitors and neurons

Starting from DIV 19, cells were seeded at 0.45 × 10^6^/cm^2^ in basal differentiation medium with 50 nM Chroman 1. After 24 hours, Chroman 1 was removed and cells were maintained in CTX media for 2 days. Media was then supplemented with 3µM CHIR99021(Wnt activator) for 10-14 days to generate hippocampal progenitors capable of differentiating into the different subregions of the hippocampus. Cells were then cultured in basal differentiation media only till DIV 40-45. Hippocampal progenitors were passaged 1:2 every 10-14 days into newly coated hLam521 plates.

For final plating, the plates or coverslips were coated with poly-L-ornithine followed by hLam521. Cells were then seeded at 2-3 × 10^5^/cm^2^ in cortex medium with 50 nM Chroman 1. Media was then switched to maturation medium containing 1:1 DMEM/F12 and Neurobasal, 1 × B27, 0.5 × N2, 0.5 × ITSA, 0.5 × NEAA, 0.5 × GlutaMAX, and 0.5 × penicillin/streptomycin, with 10 ng/mL brain-derived neurotrophic factor, BDNF (R&D Systems), 10 µM γ-secretase inhibitor DAPT, 0.25 µM *N*^6^,2′-*O*-dibutyryladenosine 3′,5′-cyclic monophosphate (Tocris Bioscience), and 200 nM ascorbic acid (Sigma-Aldrich). Cells were cultured for 3-7 days and DAPT was removed. Cells were further cultured until they were fixed with PFA for downstream experiments.

### Generation and maturation of hippocampal organoids

After the removal of CHIR99021 around DIV 33-35, cells were seeded at 50-60k per well of a 96 ultra-low attachment multiple well plate (Sigma-Aldrich-CLS7007) in CTX media with 25 nM Chroman 1. After 48 hours, Chroman 1 was removed and cells were maintained in CTX media for 14 days with media change every other 2 days. The organoids were transferred into a 10 cm petri dish with maturation medium supplemented with 10 ng/mL brain-derived neurotrophic factor, BDNF (R&D Systems), 0.25 µM *N*^6^, 2′-*O*-dibutyryladenosine 3′, 5′-cyclic monophosphate (Tocris Bioscience), 200 nM ascorbic acid (Sigma-Aldrich), and cultured on an orbital shaker (Biotek-#NBT101SRC) in the incubator at 68 rpm.

To generate hippocampal and cortical organoid (HCO), hippocampal cells were cultured till DIV 40 and cortical cells were cultured till DIV 30 and seeded at 50-60k (50-50%) per well of a 96 ultra-low attachment multiple well plate (Sigma-Aldrich-CLS7007) in CTX media with 25 nM Chroman 1. After 48 hours, Chroman 1 was removed and cultured same as HOs.

### Differentiation into hippocampal regionally specified astrocytes

Cells were harvested on DIV 19 using accutase and seeded at 5000 cells per well of a 96 ultra-low attachment multiple well plate (Sigma-Aldrich-CLS7007) in CTX media with FGF2. The plates were centrifuged at 1100 rpm for 5 minutes. Under these conditions, cells formed spheres after 24 hours and cultured for another 24 hours. Media was then changed to CTX media supplemented with 3 µM CHIR99021 for 14 days with media change every other 2 days. The spheres were then transferred into a 10cm petri dish and were maintained on an orbital shaker in the incubator. Spheres were cultured in CTX media only for 2 days before switching the media to astrodifferentation medium (ADM) for a month and manually dissociated by cutting the spheres into 4 parts. ADM was comprised of Advanced DMEM/F12 (ThermoFisher Scientific, #12634-010) supplemented with 0.2% B-27 Supplement (ThermoFisher Scientific, #17504001), 0.5% N-2 Supplement (ThermoFisher Scientific, #17502001), 0.5% GlutaMAX, 1% ANTI-ANTI (ThermoFisher Scientific, #15240-062), as well as 20 ng/ml Human LIF Protein, HEK293 (Medchem Express, #HY-P73276) and 20 ng/ml EGF (ThermoFisher Scientific, #AF-100-15). After a month of culture in human LIF, LIF was replaced with human recombinant bFGF (StemCellTech, #78003) and cultured for additional 2-3 months. Spheres were manually dissociated by cutting the spheres into 4 parts every 2-3 weeks and transferred to a new petri dish.

Astrocyte expansion was achieved by dissociating the spheres with NeuroCult™ Enzymatic Dissociation Kit for Adult CNS Tissue (Mouse and Rat) (StemCellTech, #05715) and plated on Matrigel (Corning)-coated flask at 10^6^ per T25 flask. The cells were maintained in the ADM supplemented with 20 ng/ml human recombinant bFGF and 20 ng/ml EGF (ThermoFisher Scientific, #AF-100-15). The cells were harvested at 70-80% confluency and reseeded at 0.5 × 10^5^ per T75 flask and maintained in ADM supplemented with 20 ng/ml human recombinant bFGF and 20 ng/ml EGF. To confirm the absence of neuronal contamination, cells were seeded into 96-well plates and subjected to immunostaining for MAP2 and GFAP. To ensure biological reproducibility, independent replicates were generated by maintaining separate neurospheres for at least one month prior to initiating terminal differentiation protocols.

### Construction of MPPED1-H2B-TdTomato lentiviral reporter plasmid

To generate the MPPED1-H2B-TdTomato reporter construct, a 789-bp (UCSC Genome Browser on Human (GRCh38/hg38) - chr22:43,411,019-43,411,807) genomic fragment immediately upstream of the MPPED1 transcription start site, encompassing the predicted promoter and proximal enhancer regulatory elements, was identified and validated in silico. Putative transcriptional regulatory activity was assessed using the Berkeley Drosophila Genome Project (BDGP) Neural Network Promoter Prediction tool (https://www.fruitfly.org/seq_tools/promoter.html) and Promoter 2.0 Prediction Server (https://services.healthtech.dtu.dk/services/Promoter-2.0/), both of which supported promoter functionality within the selected region.

A promoter-specific lentiviral reporter vector was subsequently engineered and packaged by VectorBuilder. The resulting construct, designated pLV[Exp]-(MPPED1 promoter)>H2B-TdTomato (Vector ID: VB260410-1055fkq), drives the expression of a nuclear-localized TdTomato fluorescent reporter under the control of the MPPED1 regulatory sequence. Lentiviral particles were produced and quality-controlled by the manufacturer, with viral titers determined using a p24 enzyme-linked immunosorbent assay (ELISA).

Frozen lentiviral particles were thawed immediately prior to use and applied to differentiated cells at a multiplicity of infection (MOI) of 40 in the presence of polybrene. After 24 hours, the medium was replaced with fresh culture medium. Cells were then maintained under standard culture conditions until fixation and downstream processing for immunofluorescence analysis.

### Immunocytochemistry, microscopy, and quantification

Immunocytochemistry was performed as previously described^93–95^ (see **Supplementary Material 1** for a list of primary and secondary antibodies). Briefly, cells were fixed for 10 minutes in 4% paraformaldehyde. Cells were permeabilised and blocked in 2% donkey serum, 0.1% Triton-X in PBS-/- with sodium azide (Blocking and permeabilisation buffer) for 20-30 minutes at room temperature. Fixed cells with primary antibodies were incubated at 4°C in the blocking and permeabilisation buffer overnight. Secondary antibodies were incubated on the fixed cells for 1-2 hours at room temperature. After PBS washes and nuclear counterstaining with Hoechst 33342, cells were mounted in Fluoromount-G™ Mounting Medium (#00-4958-02). Mounted cells were stored at 4°C before imaging. Images were captured on either an EVOS epifluorescence microscope, Operetta High Content Imaging, or a Zeiss LSM900 confocal microscope. For quantification of cell numbers and markers, images across wells were generated with at least five fields of view from multiple wells. Images were counted using the CellProfiler software. All data visualisations were generated using custom Python scripts, and statistical analyses were performed using the SciPy statistics library (Python v3.10).

For synapse quantification, images were processed using the SynBot macro as previously described^72^. Atleast 9 FOV images per independent biological replicate were converted to RGB and using the SynBot macro with the default parameters of the manual thresholding method, we quantified synapse numbers. **Supplementary Material 2** presents a list of all primary and secondary antibodies used.

### Gene expression analysis

For qPCR, cells were harvested at different stages of differentiation and centrifuged at 200 ***g*** for 5 minutes. After centrifugation, the supernatant was removed, and the cell pellet was resuspended in 100 mL of RNAlater (ThermoFisher Scientific # AM7020) and stored at −80°C. RNA was extracted using an PureLink™ RNA Mini Kit (ThermoFisher Scientific # 12183018A). After extraction, RNA was quantified using the NanoDrop 2000 UV Visible Spectrophotometer. RNA samples were then stored at –80°C or used for downstream qPCR analysis using the PrimeTime™ One-Step 4X Broad-Range Master Mix (Integrated DNA Technologies #10011746). We used 8-10 ng of RNA for each reaction using in-house designed primers and probes under the following conditions: 55°C for 10 minutes, 95°C for 1 minute, and 40 cycles of 95°C for 15 seconds and 60°C for 60 seconds. Ct values were normalised to housekeeping gene GAPDH. All gene expression data were determined using 2^−dCt^ from at least three independent biological replicates. **Supplementary Material 1** presents a list of all primers and probes used.

### RNA sequencing and quality control

Bioinformatic processing of the bulk RNA-seq samples was performed by the Australian Genome Research Facility (AGRF Bioinformatics). A total of 17 human poly(A)-selected, stranded RNA-sequencing libraries were prepared using the Illumina TruSeq Stranded mRNA protocol and sequenced as 150-bp paired-end reads on an Illumina NovaSeq X Plus platform. Primary sequence processing, including demultiplexing and sequencing quality control, was performed using the Illumina DRAGEN BCL Convert (v07.031.732.4.3.6) pipeline.

Sequencing quality was assessed according to AGRF quality control criteria. Across all 17 samples, ≥82% of bases achieved a Phred quality score of Q30 or higher. Reads were further assessed for Illumina adapter contamination, overrepresented sequences, and potential cross-species contamination. All samples met the specified quality control criteria and were retained for downstream analysis.

### Reads alignment

Following quality control, the cleaned sequencing reads were aligned against the Homo sapiens genome (Build version HG38). Read alignment was performed using STAR (v2.3.5a). Alignments were output in BAM format and indexed with SAMtools (v1.5).

### Differential expression analysis

The initial gene count matrix, comprising 39,533 genes across 17 samples, was provided by AGRF Bioinformatics **(Supplementary Material 4)**. Genes with a total count below 10 across all samples were removed, resulting in a final matrix comprising 26,293 genes across 17 samples, which was used for differential expression analysis. Differential expression analysis was performed using PyDESeq2 (v0.5.4)^96^, a Python implementation of the DESeq2 method^97^. Samples were grouped into discrete developmental stages by days in vitro (DIV 0, DIV 11, DIV 40, DIV 60, DIV 82-100, DIV 154-200). Genes with fewer than 10 total counts across all samples were removed prior to analysis. Size factors were estimated internally using the median-of-ratios method as part of the PyDESeq2 workflow, with DIV 0 as the reference level.

Differentially expressed genes were identified using Wald tests performed directly against a log fold change (LFC) of 1 with Benjamini-Hochberg-adjusted p-values <0.05 considered significant. For visualisation and ranking, LFC estimates were shrunk using the *apeglm* estimator^98^ as part of the PyDESeq2 workflow; shrinkage was applied to the fold-change estimates for effect-size visualisation and did not affect the differential-expression significance calls, which were based on the threshold-based Wald tests.

### Gene ontology enrichment analysis

Gene Ontology (GO) enrichment analysis was performed separately on the up- and down-regulated differentially expressed gene lists using the Enrichr API^99,100^ as implemented in gseapy^81^, querying the “GO_Biological_Process_2026” gene set library^101,102^. Enrichment was assessed by Fisher’s exact test, and terms were ranked by Benjamini–Hochberg adjusted p-values; terms with adjusted-p < 0.05 were considered significantly enriched. For each term, the gene ratio was calculated as the number of query genes annotated to the term divided by the total number of genes in the query list.

Simple-sample gene set enrichment analysis (ssGSEA) was performed using gseapy to calculate gene signature enrichment scores for individual bulk RNA-seq samples. The resulting enrichment scores were normalized across samples.

### Single-nucleus RNA reference data

A publicly available postnatal single-nucleus RNA-seq (snRNA-seq) dataset of human hippocampus^82^, spanning five developmental stages (infancy, childhood, adolescence, adulthood, and aging), served as the reference cell-type atlas. Raw data are available from the Gene Expression Omnibus (GEO) under accessions GSE185553, GSE185277, and GSE198323. To resolve glutamatergic neurons at finer resolution, a second adult human hippocampal snRNA-seq dataset^65^ containing refined glutamatergic neuronal subtype annotations, was used as a complementary reference. Raw data for this dataset are available from GEO under accession GSE264624, and preprocessed data were obtained through the Bioconductor ExperimentHub package (**“humanHippocampus2024“**). Raw data for this dataset are available from GEO under accession GSE264624, while preprocessed data were obtained through the Bioconductor ExperimentHub package (“humanHippocampus2024”). Additional details on retrieval of the preprocessed datasets are provided in **Supplementary Table 28**.

### Single-nucleus RNA label transfer and deconvolution

Both reference datasets had undergone quality-control filtering and batch correction as described in their respective original publications. Prior to label transfer, the two snRNA-seq datasets were restricted to their shared gene set and processed using the same workflow, including total-count normalisation, log-transformation, and scaling. Principal component analysis (PCA) was performed on the adult hippocampal reference dataset, and the cells assigned to the glutamatergic neuronal (Glut.N) cluster in the postnatal reference were projected onto the resulting 30-component PCA embedding using sc.ingest. Only the Glut.N population was subjected to label transfer; all other postnatal cell clusters retained their original broad-level annotations. This two-step annotation strategy enabled finer resolution of glutamatergic neuronal populations, including CA1, CA2, CA3, and granule cell (GC) subpopulations. The resulting annotated snRNA-seq reference comprised 224,464 nuclei and 16,254 genes across 19 cell types. **Supplementary Material 5** contains the feature, barcode, matrix, and metadata files comprising the high-resolution snRNA-seq reference dataset used for cell-type deconvolution.

### Cell type deconvolution

The finely annotated single-nucleus reference was used to estimate cell-type proportions in the bulk RNA-seq samples through reference-based deconvolution with BisqueRNA (v1.0.5)^99,100^ in R (v4.5.2). The *ReferenceBasedDecomposition* function was applied to genes shared by the single-nucleus reference and bulk datasets, generating per-sample proportion estimates for 19 cell types.

### Code Availability

**Supplementary Material 6** contains the code files for cell type deconvolution and bulk RNA-seq differential expression analysis. Other code files are available upon request.

### Data Availability

Raw FASTQ and aligned BAM files are available upon request. Pickle files containing electrophysiological and Granger-causality network metrics are provided in Supplementary Material 3.

### Multi-electrode assay (MEA)

For electrophysiological studies, monolayer hippocampal neurons and organoids (from CLV2-10 and CLV3-1) were transferred onto poly-D-Lysine (PDL)/laminin-521-coated MEA chips (Multi Channel System). MEA chips were coated with PDL and human Laminin-521 at a concentration of 50 µg/ml and 10 µg/ml respectively. Monolayer hippocampal neurons were dissociated at day 40 and plated on PDL/laminin-521-coated MEA chips at a density of 5×10^5^ cells per MEA chip. To assess the functional properties of human pluripotent stem cell (hPSC)-derived astrocytes, we co-cultured hippocampal neurons (DIV 40) with passage 8 (P+8) hippocampal astrocytes on MEA chips. Based on published data^103,104^, we cocultured the neurons and astrocytes at a ratio of 1.4:1 (neurons:astrocytes). The organoids at DIV 55 were transferred onto PDL/laminin-521-coated chips.

The MEA chips used for this study have 64 electrodes. All recordings were conducted using a CL1 system (v.0.8, Cortical Labs, Singapore) maintained at 37°C^105^. The recording of the baseline activity of MEA chips was performed weekly over a 5-minute period for 15-20 weeks after plating.

### MEA analysis

The following electrophysiological parameters were analysed: mean firing rate in Hz, max firing rate in Hz, variance of the firing in Hz², criticality analysis, power law observables, branching ratio, exponent relationship, Deviation from Criticality Coefficient (DCC), information entropy, and burst detection analysis which assessed both the number of bursts, the average number of spikes during bursts, and the percentage of bursting activity overall. The analysis for criticality analysis, power law observables, branching ratio, exponent relation, and DCC were partially adapted from previously published studies^79^.

### Criticality analysis

Avalanche analysis was used to characterise how close the network operates to a critical state. Avalanche onsets and terminations were defined based on crossings of a predefined threshold in the global network activity signal^79^. Individual avalanches may be triggered by spiking activity from any subset of neurons within the region of interest. For each avalanche, both the total number of spikes involved(S) and the overall event duration (D) were quantified. To assess the proximity of the cultured neuronal networks to criticality, we examined the presence of several established dynamical signatures in the data: (1) power-law behaviour of avalanche observables; (2) consistency of scaling exponent relationships; (3) the value of the branching ratio; and (4) the existence of an appropriate scaling function. These indicators constitute necessary conditions for critical dynamics and jointly satisfying them provides strong evidence that the system operates near a critical point^79,106^.

### Power law observables

A system operating at criticality consists of interacting elements, in this case the recorded electrodes, that exhibit ongoing fluctuations in activity while retaining a degree of coordination across individual units, such as correlated spiking. Critical dynamics are characterised by scale-invariant behaviour, whereby events across both temporal and spatial scales follow power-law statistics. In the networks studied here, such events correspond to continuous cascades of spiking activity, as opposed to isolated local bursts or globally synchronous network-wide discharges. These cascades are commonly referred to as *neuronal avalanches*. To examine this phenomenon in our recordings, we analysed binarized spike trains from each electrode. Each recording session was segmented into temporal bins of 50 ms, and network activity at each timestep was defined as the sum of spiking activity across all neurons within that bin. An activity threshold was then set at 40% of the median network activity computed across all-time bins. Avalanche initiation and termination were identified as the moments when the network activity crossed this threshold from below and subsequently returned below it^109^. Importantly, the results were stable across a broad range of threshold values spanning 30% to 70% of the median activity. The size of an avalanche, S, is the total number of spikes during the avalanche. The avalanche duration, D, is the time between threshold crossings. Following the approach in^110^, a truncated power-law model was fitted to the avalanche size distribution using maximum likelihood estimation as below:

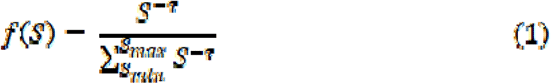

Here, denotes the power law exponent associated with the avalanche size distribution. For a given neuronal recording session in which avalanches are identified, the estimation of this model follows an iterative fitting process as described in ^111^, as outlined below:

1. Find the maximum observed avalanche size *Smax.*
2. Evaluate the three different power law exponents, *τ*, for the 3 smallest avalanche sizes observed, *S_min_*.
3. Calculate the *Kolmogorov-Smirnov* (KS) to assess the goodness-of-fit between the fitted power-law model and the empirical distribution.
4. Select the minimum KS value among all estimates, along with the corresponding *τ* and *S_min_* values.
5. Complete the estimation if 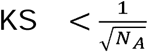 otherwise repeat steps 2 to 5 with *S_max_* reduced by 1 until this condition is met.

Steps 3–5 ensure that the observed data distribution is consistent with a power-law model rather than alternative heavy-tailed candidates, such as log-normal or stretched-exponential distributions ^111^.

The same analysis was then carried out for the set of avalanche durations, yielding the corresponding power-law exponent α for the distribution of avalanche durations.

### Exponent relation and Deviation from Criticality Coefficient (DCC)

In systems operating near criticality, an additional scaling relationship links the power-law exponents α and τ to the exponent governing the mean avalanche size (〈S〉), as a function of avalanche duration *D*^112^. We first estimated the third scaling exponent of the system, *β*, directly from the experimental data using linear regression, assuming the following exponent relationship holds for a system at criticality:

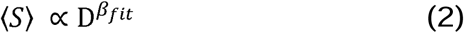

This third scaling exponent further connects the avalanche size and duration distributions and is theoretically given by:

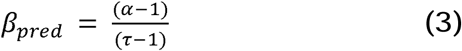

Comparing the fitted value from the empirical data (*β_fit_*) and its estimation using *α* and *τ* exponents, (*β_pred_*), a new measure is derived to evaluate the *Deviation from Criticality Coefficient* (DCC), parameterised as dcc:

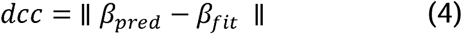

where, (*β_pred_*) and (*β_fit_*)are the predicted and fitted values of, *β* respectively.

Accordingly, lower DCC values correspond to a closer agreement between the fitted power-law model and the empirical distribution.

#### Information Entropy

A binary vector was used to represent the presence or absence of a spike in each 50 ms time bin for each recorded electrode. We then calculated the binary entropy function for every electrode using their binary spike trains.

The binary entropy function, denoted *H*(*p*) or *H_b_*(*p*) is defined as the entropy of a Bernoulli process with probability p of one of two outcomes.

Given that:

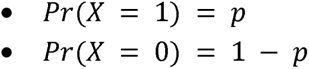

The entropy of X (measured in shannons) is calculated as:

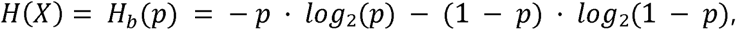

where 0 · log₂(0) is defined to be 0.

Hence, we calculated the local entropy each electrode in each time bin and averaged across time and electrodes to compare the groups of in vitro cultures.

### Burst detection analysis

To identify bursting activity, we analysed spike trains from each electrode. A burst was defined as a sequence of at least *N* spikes (default: 5) in which the inter-spike interval (ISI) between consecutive spikes did not exceed a fixed threshold (default: 1 ms). For each electrode, spike times were first sorted, and bursts were detected by scanning for sequences of spikes that satisfied both the minimum spike count and maximum ISI criteria. For each detected burst, we recorded its duration (time between the first and last spike, in seconds) and size (total number of spikes). These burst metrics were stored for downstream analysis.

### Construction of causal connectivity graphs using granger causality

To infer directed causal relationships among activities of recorded electrodes, we constructed causal connectivity graphs from spike train recordings using Granger causality analysis^113^. Recordings were obtained from 64-channel multi-electrode arrays (MEAs) sampled at 25 kHz. After excluding the reference electrodes (electrode IDs 0, 4, 7, 56, and 63), a total of 59 active electrodes were retained for analysis.

Spike times were first extracted from each recording and converted into binary spike matrices (“spikewords”) representing binned firing activity for each channel. The bin length for this analysis was set to 1 s. Each matrix entry indicated whether a spike occurred in a given bin for a given electrode. Electrodes with mean firing rates below 0.05 Hz were excluded to reduce noise from inactive channels.

Pairwise Granger causality tests were then computed between all remaining electrodes to estimate directional dependencies in spiking activity. For each ordered electrode pair (*i,j*), the corresponding time series were concatenated into a two-column matrix [*x_i_,x_j_*], and Granger causality tests were performed across lags from 1 to 50 using the F-test formulation. The minimum *p*-value across lags was used to determine significance (*p* < 0.05), and the corresponding F-statistic was recorded as the causal connection strength. Connections failing to meet this significance criterion were omitted.

The resulting weighted, directed adjacency matrix *A* ∈ ℝ^{64x64}^ was constructed such that *A*_{*ij*}_ represents the causal influence from electrode *j* to electrode *i*. Self-connections were excluded by enforcing *A*_{*ij*}_ = 0. The number of incoming and outgoing connections was also computed for each electrode to characterise its causal in-degree and out-degree.

To quantify the topological and dynamical organization of these causal networks, several standard graph-theoretical measures were computed from each adjacency matrix.

- **The total number of edges** (*E*) was calculated as the number of significant directed causal links detected in the network. average edge weight (*w̅*) was obtained as the mean F-statistic across all significant connections:

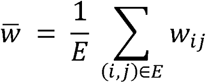
- **The clustering coefficient** was computed using the weighted variant of the Watts–Strogatz formulation^117^ as extended by Onnela et al.^118^. For an undirected and weighted graph *G* = (*V*, *E*) with edge weights *w_ij_* ≥ 0, the local weighted clustering coefficient 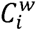 for a node is defined as:

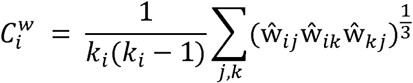 Where *k_i_* is the degree of node *i* (number of neighbors), 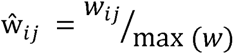 is the normalised edge weight, and 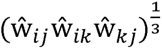 represents the geometric mean of the weights forming a triangle between nodes *i,j*, and *k*. The global clustering coefficient for the entire network is then computed as:

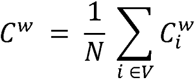 This metric ranges from 0 (no clustering) to 1 (perfect clustering), providing a normalised measure of how locally interconnected the network is. A higher *C^W^*, indicates that groups of electrodes tend to form tightly connected subnetworks, suggesting strong local synchronisation or recurrent information flow among nearby neurons, whereas lower values imply more random or sparse local interactions.
- **The mean betweenness centrality** was calculated as the average of the weighted betweenness scores across all nodes Mathematically, the betweenness centrality *C_B_*(*v*) of a node V is defined as:

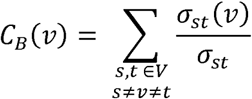 where *σ_st_* is the total number of shortest paths between nodes *s* and *t*, and *σ_st_*(*v*) is the number of those paths that pass through node *v*. For weighted networks, edge lengths are defined as the inverse of their weights (*l_ij_* = 1/*w_ij_*), such that stronger causal connections corresponded to shorter path distances. The mean betweenness centrality across all *N* = |*V*| nodes were computed as:

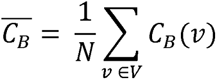 This measure captures how strongly specific electrodes act as communication hubs that mediate information flow between otherwise disconnected neuronal populations. A high mean betweenness centrality therefore indicates a more centralised network organisation dominated by a few influential nodes that integrate and redistribute signals, whereas lower values suggest a more distributed and parallel mode of information transfer across the culture.

### Statistical Analysis (Group Comparisons)

To examine statistical differences between cell lines, qPCR measurements were analysed by two-way ANOVA, followed by pairwise comparisons using two-tailed Welch’s t-tests. To avoid pseudoreplication, immunocytochemistry measurements were first averaged within each biological replicate, and cell lines were compared using a two-tailed Welch’s t-test on the biological replicate means^116^. Beeswarm plots display both the individual technical replicate measurements and the mean of the technical replicates for the immunocytochemistry measurements for each biological replicate.

Multiple-comparison correction methods were applied as appropriate, with the specific methods described in the corresponding figure captions or supplementary tables. Group values for quantitative PCR and immunocytochemistry measurements are presented as mean ± standard error of the mean (SEM).

Sensitivity analyses were performed by excluding outliers identified using the interquartile range (IQR) method within each group. Outlier removal was applied only to the sensitivity analyses and not to the primary analyses. For each metric, group distributions were visualised using violin and raincloud plots. For each metric, group distributions were visualised using violin and boxplots. Statistical differences between groups were assessed using Kruskal-wallis, followed by a Dunn’s post-hoc test to account for unequal sample sizes. Significance thresholds were annotated as follows: *: p < 0.05, **: p < 0.01, and ***: p < 0.001.

### Survival Analysis

To compare the time-to-event distributions between electrophysiological and functional connectivity metrics, the log-rank (Mantel–Cox) test was used to assess statistical differences in survival curves, which were visualized using Kaplan–Meier plots. Statistical significance was defined as p < 0.05.

## Supporting information

Supplementary Data - Electrophysiological Data (Fig 5-7)

Supplementary Data- Immunochemistry & Quantitative PCR Data

Supplementary Material-Antibody List

Supplementary Material-qPCR Primers

Supplementary Table 27 - Bulk Samples Metadata

## Supplementary Data and Methods

Supplementary Figures 1–5 are included below for convenience; high-resolution images are available to download.

Supplementary Tables 1–26 contain the full results of all statistical analyses and are available to download as a separate file.

Supplementary Table 27 provides metadata for the bulk RNA-seq samples, and Supplementary Table 28 describes the snRNA-seq datasets used.

Supplementary Table 29 contains gene expression values for all bulk RNA-seq samples.

Supplementary Tables 30 and 31 list the top 500 upregulated and downregulated differentially expressed genes (DEGs), respectively.

Supplementary Tables 32-46 contain DEGs from pairwise comparisons between all culture stages.

Supplementary Tables 47-51 contain gene ontology analyses of upregulated and downregulated genes for comparisons between DIV 0 and each subsequent culture stage.

Supplementary Tables 52 and 53 contain gene expression values and the percentage of cells expressing each gene for each cell type in the snRNA-seq reference dataset used for deconvolution. Supplementary Table 54 shows the estimated cell-type proportions following deconvolution.

Supplementary Tables 55-68 provide the raw data needed to reproduce the immunostaining analyses figures for Figure 1-3.

Supplementary Methods are available to download as a separate file.

## Acknowledgements

This work was funded by Cortical Labs Pty Ltd. The authors gratefully acknowledge the Biological Optical Microscopy Platform, University of Melbourne for their support & assistance in this work. The authors also thank Professor Edouard G. Stanley for the RM3.5 TdTomato hPSC cell line and Dr. Nicole Kerlero de Rosbo and Dr Sharmony Kelly for their invaluable assistance in proofreading the final manuscript.

## Author contributions

K.D.A.B and B.J.K conceived and designed the experiments. K.D.A.B and C.D performed neuronal differentiation and other cellular and molecular assays. Data analysis and interpretation were carried out by K.D.A.B., C.D. and H.W.C. Electrophysiological recordings and data acquisition were conducted by K.D.A.B. and C.D., while electrophysiological data analysis was performed by F.H. and H.W.C. iPSC lines were generated by B.W. The manuscript was written and revised by K.D.A.B., C.D., F.H., H.W.C., B.W., M.D., and B.J.K. The work was supervised by B.J.K. All authors discussed, commented and agreed on the manuscript.

## Supplementary Figures

Supplementary Figure 1:

A- Immunostaining analysis of SOX2+ progenitors and TBR2+ IPCs at DIV 22 for two hPSC lines. Scale bar is 100 µm.

B- Immunostaining analysis of SOX2+ progenitors and ZO-I+ lumens showing the gradual increase in diameter of the lumen for two hPSC lines. Scale bar is 100 µm and 50 µm.

Supplementary Figure 2:

A- qRT-PCR analysis confirming the increased expression of neuronal markers and reduced expression of pluripotency markers for two hPSC lines (male and female, n = 3 independent experiments each, data are mean ± SEM).

B- Quantification for PROX1+ and TBR2+ cell populations to validate the optimised CHIR99021 concentration at DIV 40 for two hPSC lines (male and female, n = 3 independent experiments each). The Y axis refers to the % of cells expressing the marker over the total of cells counted as per Hoechst staining. * *p* < 0.05, ** *p* < 0.01, \*\*\**p* < 0.001

Supplementary Figure 3:

A- Representative images of FOXG1+ and MAP2+ neurons at DIV 60. Scale bar is 100 µm.

B-Quantification of CTIP2+, MAP2+, PROX1+, ELAVL2+, SATB2+, STMN1+, CALB+ and WFS1+ cell populations at DIV 60 for two hPSC lines (male and female, n = 3 independent experiments each, data are mean ± SEM). The Y-axis refers to the % of cells expressing the marker over the total of cells counted as per Hoechst staining.

C- Immunostaining analysis of TAU+, ELAVL2+, PROX1+, OCT6, WFS1+ and CTIP2+ neurons showing large somata pyramidal neurons and small somata neurons at DIV 90 for the female cell line. Scale bar is 50 µm

D- Immunostaining analysis of TAU+, ELAVL2+, PROX1+, OCT6, WFS1+ and CTIP2+ neurons showing large somata pyramidal neurons and small somata neurons at DIV 90 for the male cell line. Scale bar is 50 µm

E- Immunostaining of DIV 90 cultures demonstrating heterogeneous CA3 neuronal populations defined by SCGN, ELAVL2, and GRIK4 expression for two hPSC lines. Scale bar is 50 and 100 µm.

F- Representative brightfield images of generated HOs at two different ages, DIV 200 (right panel) and DIV 400 (left panel). Morphometric analysis of the degree of circularity, roundness and solidity of the organoids across various cell lines (CLV4-4 (female), CLV5-1 (female), CLV3-1 (male) and RM3-5 Tdtomato (male), n = 3 independent cell lines). Scale bar is 500 µm.

Supplementary Figure 4:

A - Changes over time in electrophysiological metrics from months 2–14 in vitro. Lines and points show the mean across cultures per month, and shading shows ± 1 SD.

B - Violin plots of total spike count, maximum firing rate (Hz), and burst frequency for hippocampal culture group. Each violin shows the distribution of values across recordings, with culture-level means indicated. Statistical comparisons between groups were performed on culture-level means using the Kruskal–Wallis test followed by Dunn’s post hoc test with Sidak correction; *p < 0.05, **p < 0.01, ***p < 0.001.

C - Kaplan–Meier survival curves showing culture longevity as a function of meeting predefined functional activity thresholds for each metric shown in (C). Survival probability is plotted against days in vitro (DIV) for each culture group. Thresholds: total spike count ≥ 500, maximum firing rate (Hz) ≥ 50, burst frequency ≥ 0.007. Cultures failing to meet the specified threshold were considered to have failed the criterion. Shaded regions indicate 95% confidence intervals.

D - Violin plots of total spike count, maximum firing rate (Hz), and burst frequency for hippocampal culture group upon exclusion of outliers with IQR. Each violin shows the distribution of values across recordings, with culture-level means indicated. Statistical comparisons between groups were performed on culture-level means using the Kruskal–Wallis test followed by Dunn’s post hoc test with Sidak correction; *p < 0.05, **p < 0.01, ***p < 0.001.

E – Raincloud plots of Granger-causality based network metrics and mean information entropy for hippocampal culture groups: total edge counts, average edge weights, clustering coefficient, betweenness centrality and mean information entropy, upon exclusion of outliers with IQR. Each raincloud shows the distribution of values across recordings, with culture-level means indicated. Statistical comparisons between groups were performed on culture-level means using the Kruskal–Wallis test followed by Dunn’s post hoc test with Bonferroni correction]; *p < 0.05, **p < 0.01, ***p < 0.001.

F- Immunostaining of DIV 90 cultures showing hippocampal neurons (NF-H, POU3F1/OCT6) and astrocytes (S100B) in two hPSC lines. Scale bar is 100 µm.

Supplementary Figure 5:

A- Volcano plots showing differential gene expression between DIV 0 and DIV 40, DIV 60, and DIV 82–100. Genes significantly upregulated at later culture stages are shown in red, significantly downregulated genes in blue, and non-significant genes in grey (adjusted p < 0.05 and |log₂ fold change| > 1). For each direction, the top 5 significant genes by adjusted p-value and the top 5 by log₂ fold change are labelled.

B - Top 10 enriched Gene Ontology (GO) biological processes among genes upregulated and downregulated at DIV 154–200. Gene ratio is plotted on the x-axis, point size indicates the number of genes in each term, and point colour indicates the adjusted p-value.

C - Dot plot showing the expression of representative marker genes across cell types in the adult hippocampal snRNA-seq atlas and the postnatal hippocampal snRNA-seq dataset used to generate the reference snRNA-seq dataset for cell type deconvolution. Marker genes shown are RELN (Cajal–Retzius); CNGB1 and PROX1 (GC); CLMP, COL5A2, FIBCD1 and MPPED1 (CA1); EYS and RGS14 (CA2/MC); KIT, GRIK4, TSPAN18, AMPH, NECTIN3, TRPS1, CHRM5 and ELAVL2 (CA3.1 and CA3.2); GAD1 and GAD2 (GABA.N); FREM3 and SOX11 (HATA/AHi); CBLN2 and CUX2 (L2/3); SATB2 (CA1 and L2/3); TSHZ2 (ProS/Sub, L2/3 and HATA/AHi); RORB (L5); FOXP2 and TLE4 (L6/6b); CNTN6, COL24A1, NTNG1, TOX (ProS/Sub); LHFPL3 and PDGFRA (OPC); MBP and MOBP (oligodendrocytes); AQP4, GFAP and SLC1A2 (astrocytes); FLT1 (endothelial); APBB1IP (microglia); CLIC6 and TTR (choroid plexus); and DNAH9 (ependymal). Note that WFS1 is more highly expressed in the ProS/Sub cluster than in CA1 in the adult hippocampal snRNA-seq atlas. Dot size shows the fraction of cells in each group that express the gene, and colour shows scaled mean expression.

D – UMAP of the reference single-nucleus RNA-seq dataset of the human hippocampus used for deconvolution, coloured by annotated cell type and shown separately by developmental stage.

Supplementary Figure 6:

A- UMAP feature plots of selected representative marker genes, grouped by the hippocampal subregion each marker represents.

Supplementary Figure 7:

A- Heatmap showing z-scored expression of genes enriched in the CA1, CA3 and DG neuronal population. Marker selection was based on enriched CA1, CA3 and DG genes previously identified. Genes are hierarchically clustered.

B- Dot plot illustrating the expression of representative CA1, CA3 and DG marker genes across cell types in a human hippocampal snRNA-seq reference dataset.

