## Supplementary figures and images for "Coordinated tri-inhibition generates human CA1, CA3 and DG neurons and hippocampal astrocytes forming long-lived and electrophysiologically complex cultures"

### Supplementary Figure 1

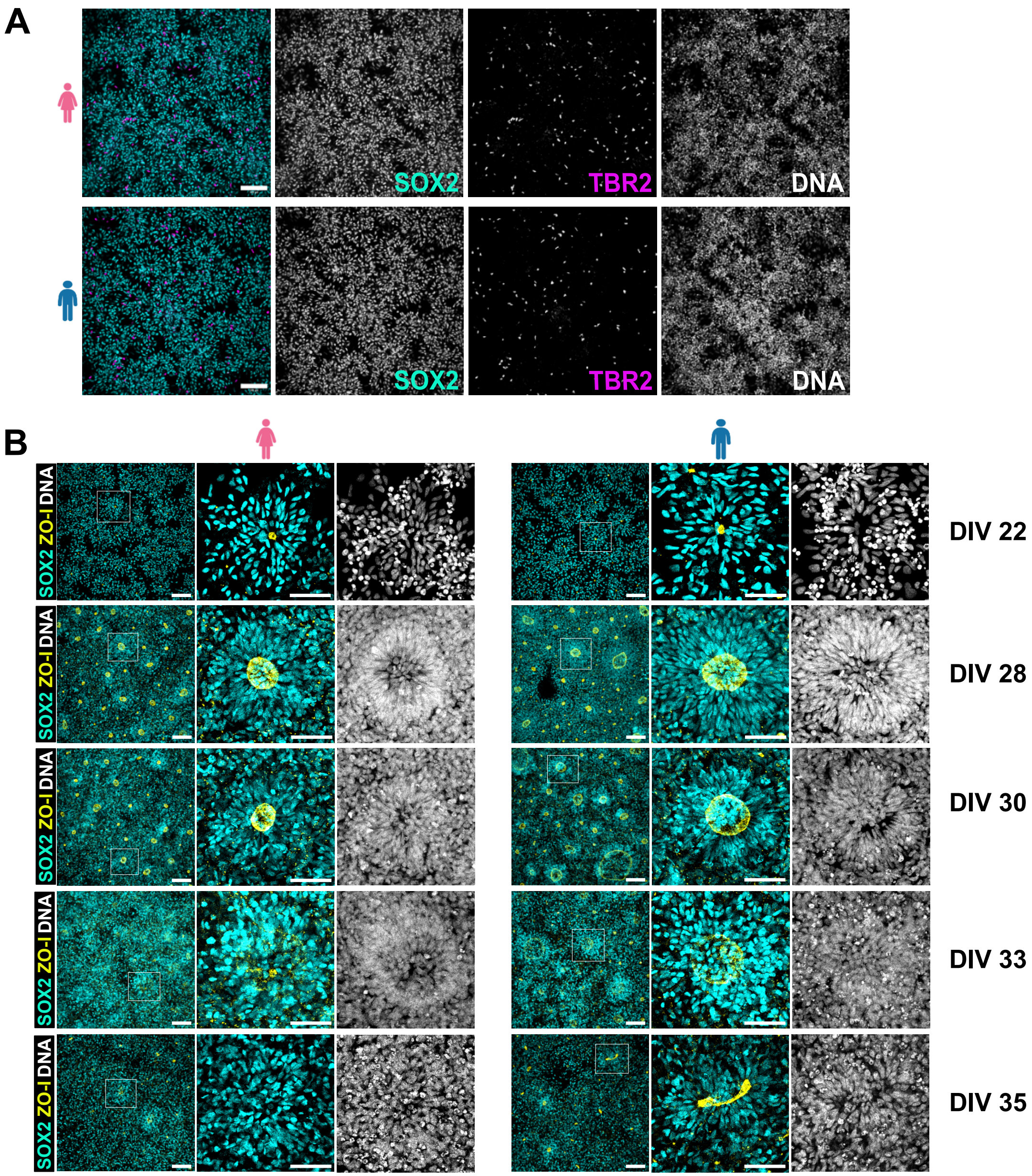

### Supplementary Figure 2

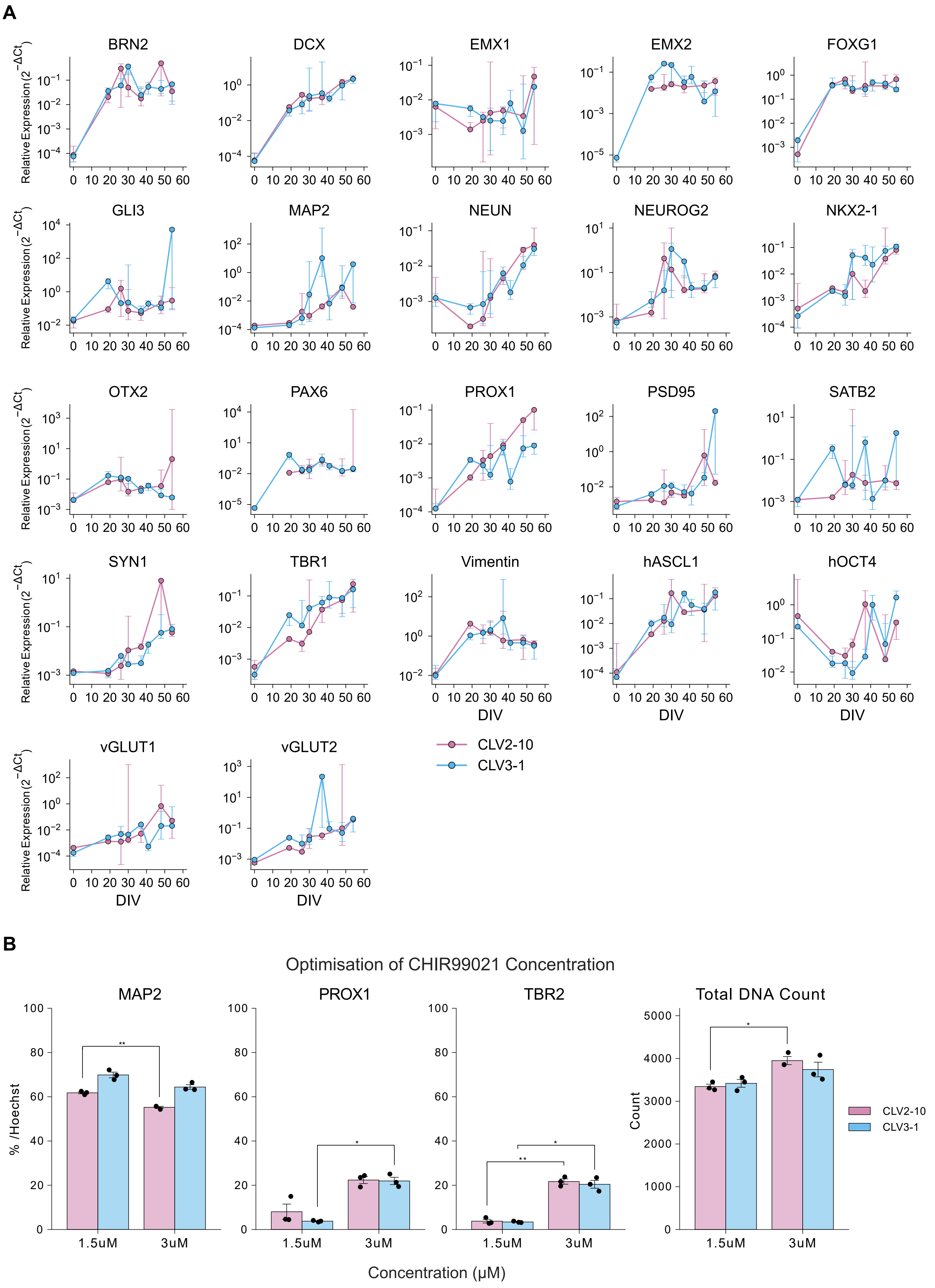

### Supplementary Figure 3

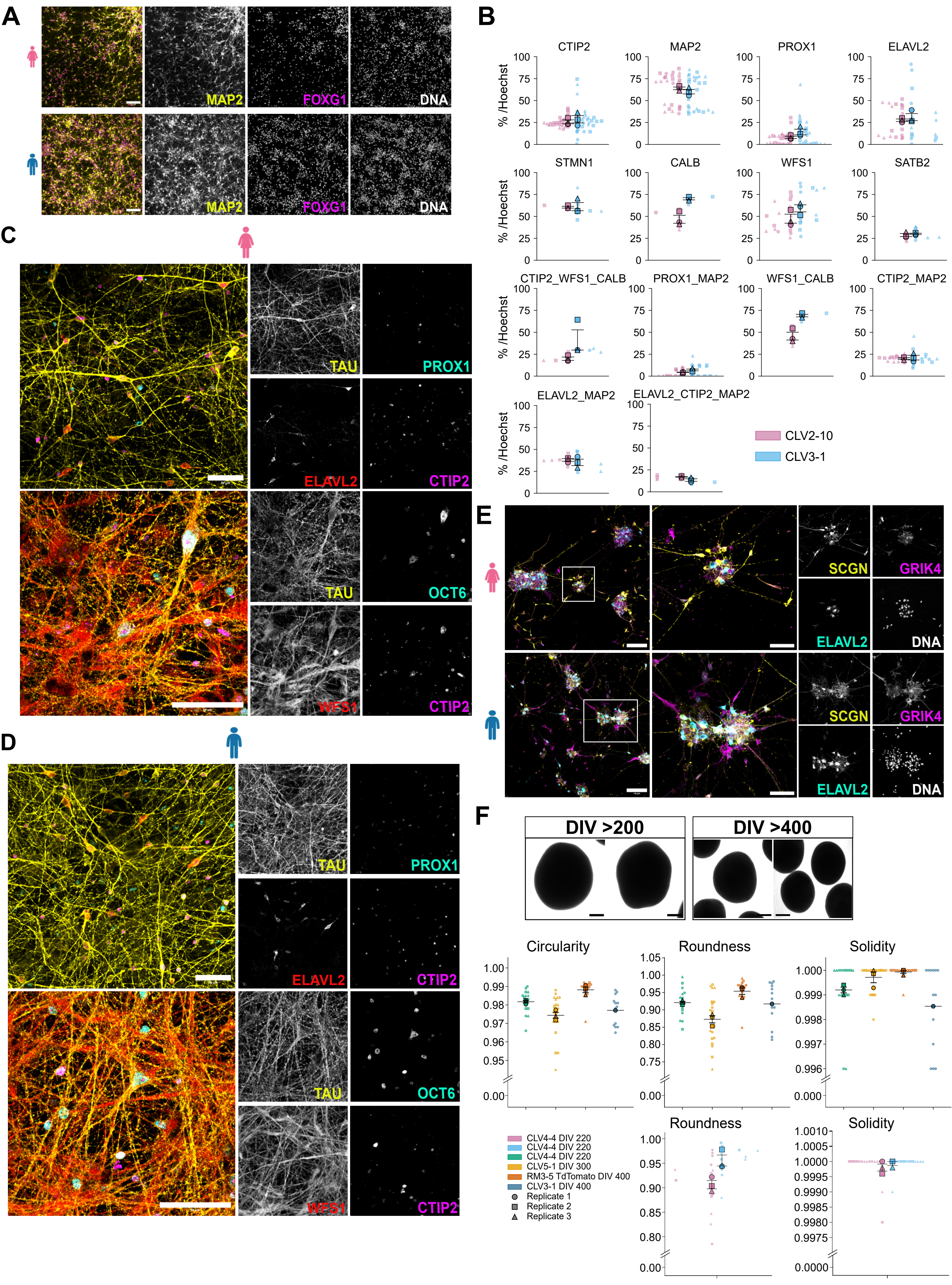

### Supplementary Figure 4

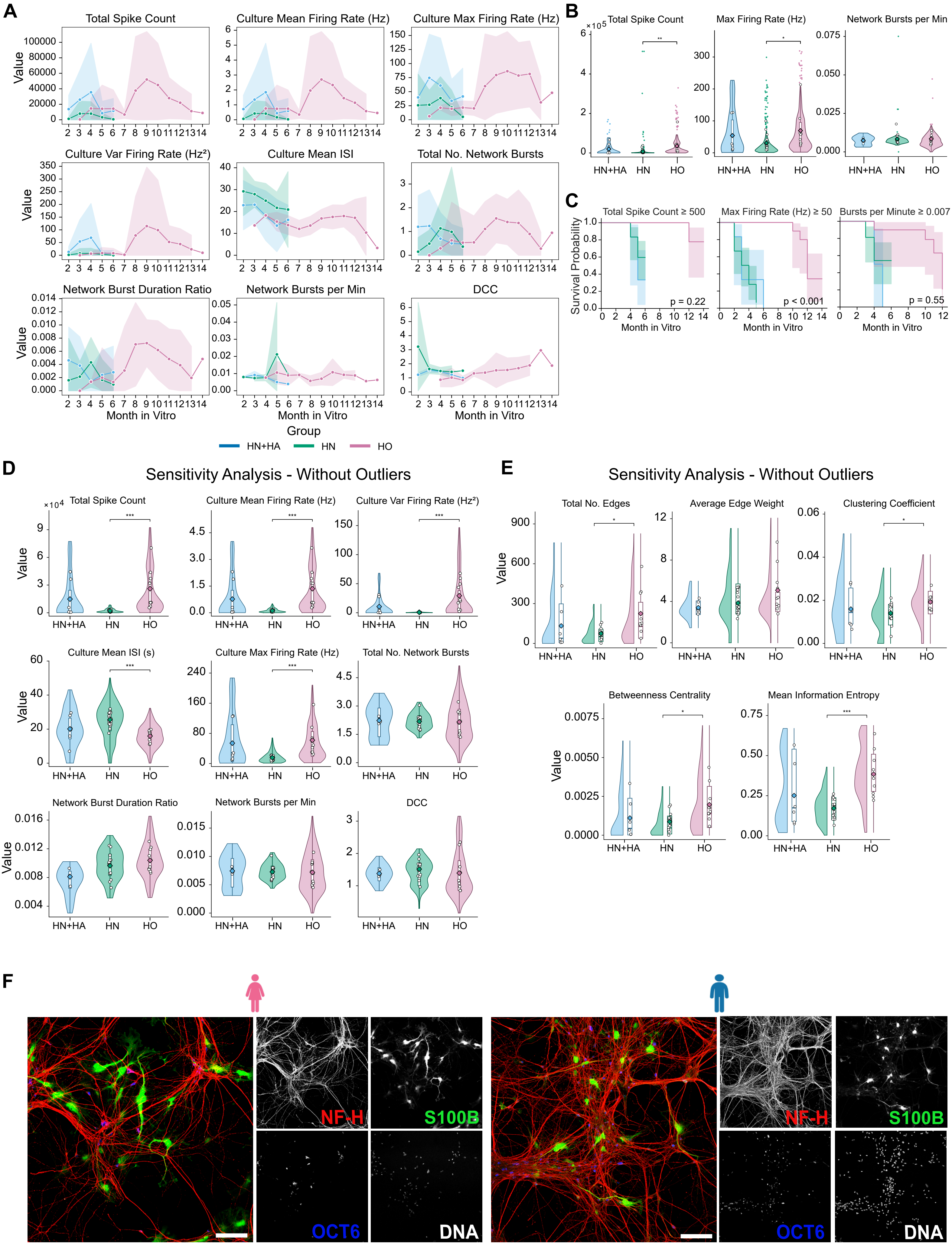

### Supplementary Figure 5

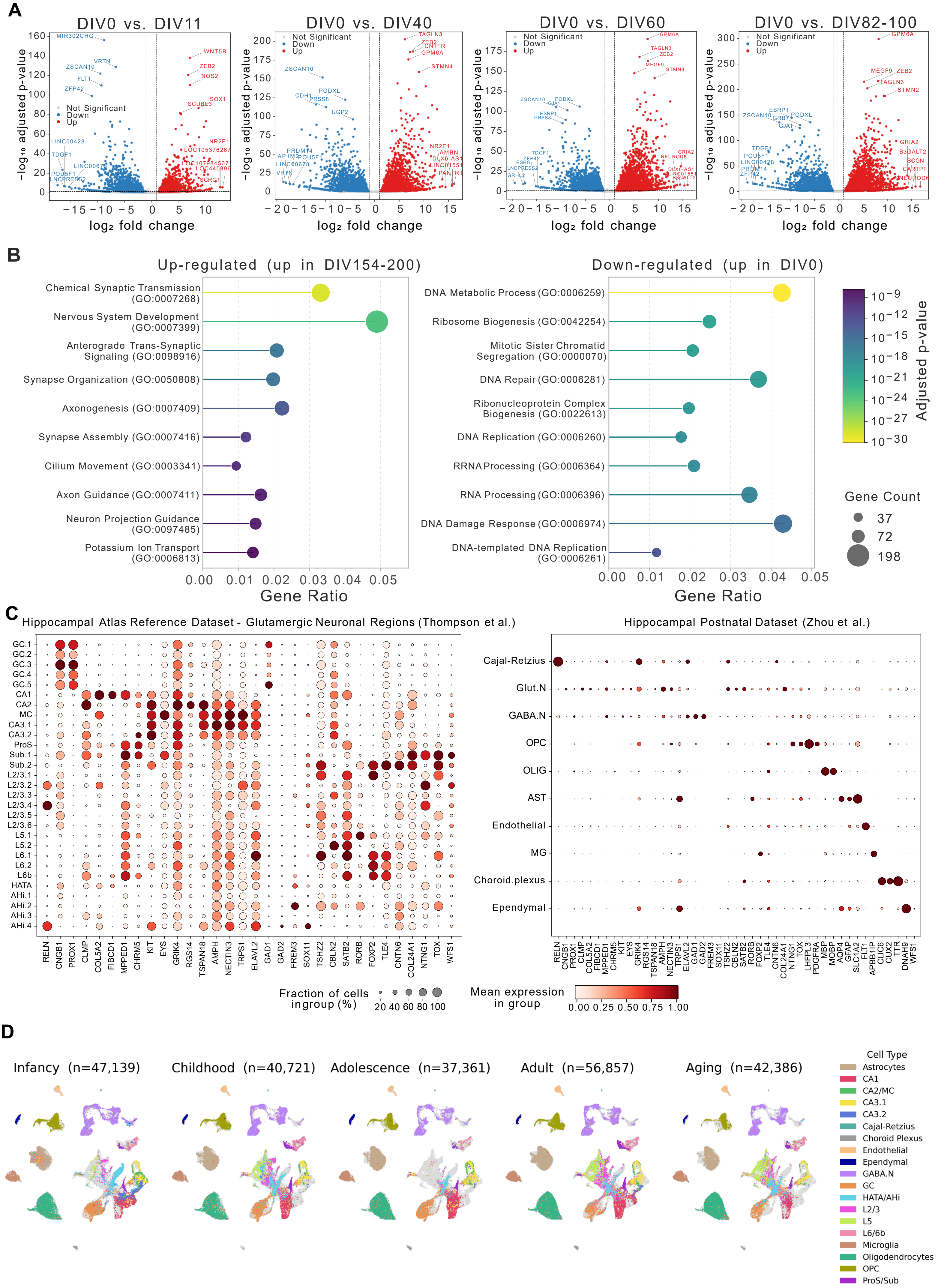

### Supplementary Figure 6

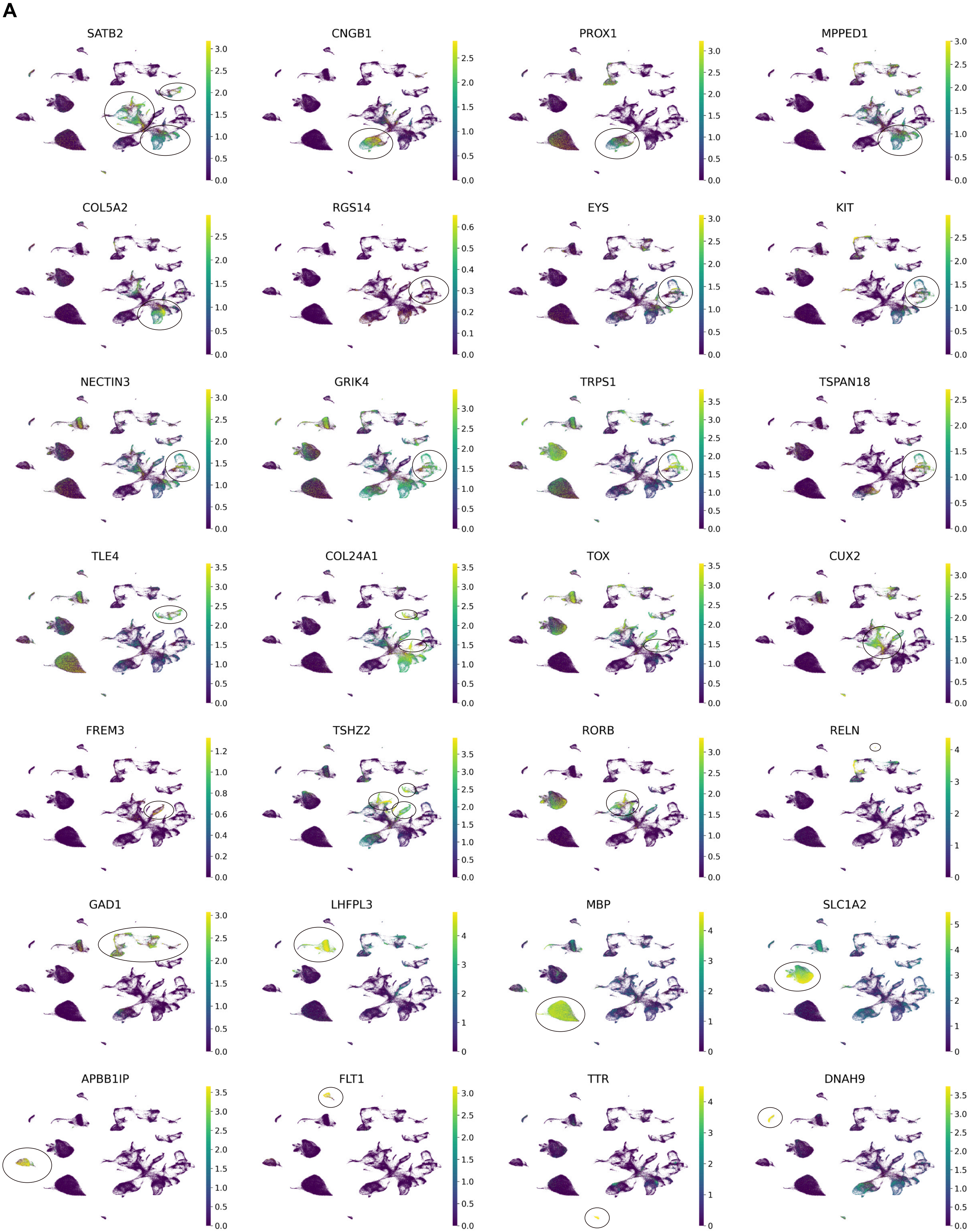

### Supplementary Figure 7

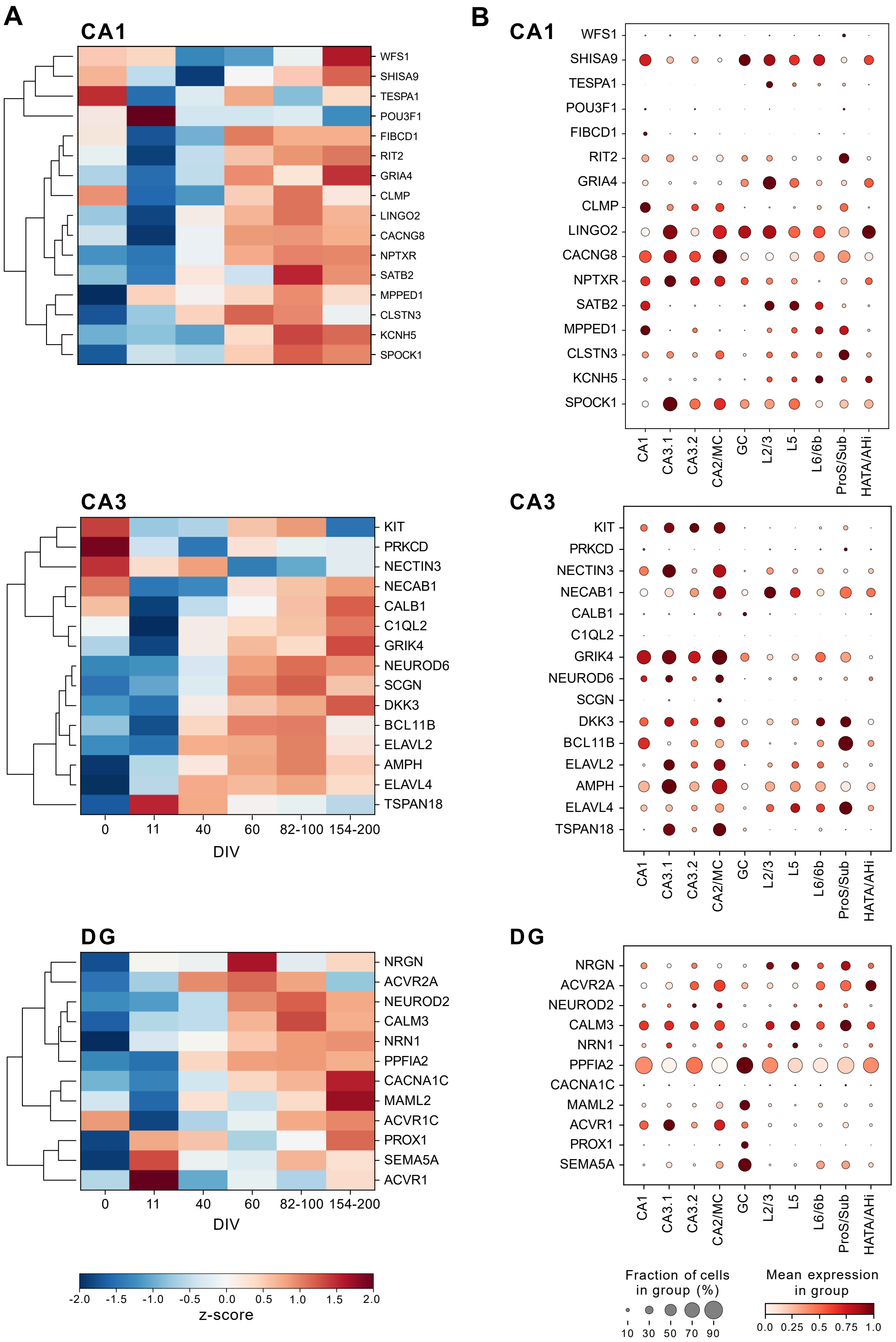
